# Molecular surveillance of multiplicity of infection, haplotype frequencies, and prevalence in infectious diseases

**DOI:** 10.1101/2025.03.13.642971

**Authors:** Henri Christian Junior Tsoungui Obama, Kristan Alexander Schneider

## Abstract

**Background:** The presence of multiple different pathogen variants within the same infection, referred to as multiplicity of infection (MOI), confounds molecular disease surveillance in diseases such as malaria. Specifically, if molecular/genetic assays yield unphased data, MOI causes ambiguity concerning pathogen haplotypes. Hence, statistical models are required to infer haplotype frequencies and MOI from ambiguous data. Such methods must apply to a general genetic architecture, when aiming to condition secondary analyses, e.g., population genetic measures such as heterozygosity or linkage disequilibrium, on the background of variants of interest, e.g., drug-resistance associated haplotypes.

**Methods and Findings:** Here, a statistical method to estimate MOI and pathogen haplotype frequencies, assuming a general genetic architecture, is introduced. The statistical model is formulated and the relation between haplotype frequency, prevalence and MOI is explained. Because no closed solution exists for the maximum-likelihood estimate, the expectation-maximization (EM) algorithm is used to derive the maximum-likelihood estimate. The asymptotic variance of the estimator (inverse Fisher information) is derived. This yields a lower bound for the variance of the estimated model parameters (Cramér-Rao lower bound; CRLB). By numerical simulations, it is shown that the bias of the estimator decrease with sample size, and that its covariance is well approximated by the inverse Fisher information, suggesting that the estimator is asymptotically unbiased and efficient. Application of the method is exemplified by analyzing an empirical dataset from Cameroon concerning anti-malarial drug resistance. It is shown how the method can be utilized to derive population genetic measures associated with haplotypes of interest.

**Conclusion:** The proposed method has desirable statistical properties and is adequate for handling molecular consisting of moderate number of multiallelic molecular markers. The EM-algorithm provides a stable iteration to numerically calculate the maximum-likelihood estimates. An efficient implementation of the algorithm alongside a detailed documentation is provided as supplementary material.

**Author summary:** Malaria annually causes 263 million infections and 596,000 deaths. Control efforts are challenged by factors like spreading drug resistance. Monitoring pathogen variants at the genetic level (molecular surveillance), especially those linked to drug resistance, is a public health priority. A major challenge is the presence of multiple, genetically distinct pathogen variants (characterized by several genetic markers) within infections (multiplicity of infection). Because genetic assays do not provide phased information in this context, ambiguity in reconstructing the actual variants present in an infection arises. This challenge is not limited to malaria. Probabilistic methods are required to phase genetic data, i.e., to reconstruct the pathogen variants present in infections. As such, we introduce a statistical method to estimate the distribution of pathogen variants at the population level from unphased molecular data obtained from disease-positive specimens. This is a combinatorially difficult task, as the number of possible genetic variants grows exponentially with the amount of genetic information included. Although the method applies to data with an arbitrary genetic architecture, its application is constrained by computational limitations. The method’s adequacy is explored and used to analyze a malaria dataset from Cameroon to guide applications. A stable numerical implementation is provided.

## Introduction

Epidemiological surveillance of infectious diseases is increasingly augmented by molecular/genetic methods. This facilitates a shift from symptoms-based to pathogen-specific diagnostics and allows monitoring of specific pathogen variants of interest, such as variants associated with antimicrobial resistance, and their routes of transmission. Molecular surveillance became popular for a variety of pathogens of interest [1], due to advances in molecular/genetic methods.

Pathogen variants are typically characterized by their allelic configuration at certain loci/markers, e.g. by SNP barcodes, a microsatellite (STR) profile, or micro-haplotypes. The presence of several pathogen variants within an infection is common in many diseases. For instance, in malaria infections, it is well-recognized that distinct pathogen variants can be co-transmitted (by the same or different mosquitoes), a phenomenon commonly referred to as complexity of infection (COI) or multiplicity of infection (MOI) [2]. Especially in malaria, MOI is recognized, because it scales with transmission intensities, however not necessarily in a linear way [3, 4]. Additionally, MOI in malaria has far-reaching implications, as it mediates the amount of recombination between parasite variants and thereby changes the population genetics underlying malaria evolution, e.g., with regard to drug resistance [5]. Due to MOI in malaria, it is also important to distinguish between a variant’s frequency (its relative abundance) and its prevalence (the probability of observing it in an infection) [2]. The latter is of clinical and epidemiological relevance, the former is of relevance for molecular surveillance, e.g., when studying the spread of drug resistance.

MOI is unfortunately ambiguously defined in the literature, with discrepancies between verbal and formal definitions, and the actual quantity referred to as MOI. These differences are discussed in detail in [2]. Namely, MOI is formally defined in most statistical models as the number of super-infections with the same or different pathogen variants, assuming that exactly one variant is transmitted at each infective event biased (cf. [2]). In the following, this definition of MOI is used. Verbally, MOI is often referred to as the number of different pathogen variants within an infection, which is a derived quantity from the formal definition of MOI. Note, that co-transmission of different pathogen variants (co-infections) is ignored in the formal definition of MOI. Estimating MOI is typically coupled with estimating the frequency spectra of pathogen variants. Several methods have been proposed, often in the context of malaria, although these methods are not specific to this disease. Ad-hoc methods, e.g., [6, 7], are straightforward to apply but typically biased (cf. [2] for a detailed discussion). Alternative methods relying on formal statistical frameworks are more or less based on the same probabilistic model (cf. [2]). Differences occur in: (i) the quantities to be estimated (whether MOI or a derived parameter is estimated; cf. [2]); (ii) the genetic architecture allowed by these approaches, e.g., single marker [8, 9]), two multiallelic markers [8, 10], a set of biallelic markers [11, 12], or multiple mutiallelic markers [13, 14]; (iii) whether MOI is estimated explicitly [15, 16]; (iv) whether the probabilistic model uses approximations [11, 17, 18]; (v) the assumptions regarding the underlying distribution of MOI [2, 19]; (vi) whether heuristic plug-in estimates are required for some parameters [20]; (vii) whether bias-corrections are applied [21]); (viii) the way missing data is handled [22]; and (ix) whether a Bayesian (e.g., [11, 14, 16]) or a frequentist approach (e.g., [8, 9, 12]) is being used.

Estimators for MOI and variant frequencies typically do not have an explicit form and have to be calculated numerically. In the frequentist case, this is often achieved by using the EM-algorithm [10, 12, 17–19, 22–24]. Importantly, the asymptotic properties of the estimators were studied in detail for some methods – mainly in the simplest cases assuming a genetic architecture consisting of a single marker [21, 22, 25]. In these cases it can be shown that the probabilistic models fall into the class of exponential families, proving the existence, uniqueness, asymptotic unbiasedness, efficiency, and consistency of the estimators. Such investigations were complemented by numerical simulations to study the finite-sample properties. The same approach was used for more complicated underlying genetic architectures [10, 12]. For the simple genetic architectures, it was also shown that the maximum likelihood estimates of MOI and variant frequencies coincide with the moment estimator for MOI and variants prevalence [25]. Notably, Bayesian and maximum-likelihood estimators should be in agreement if an uninformative prior distribution of model parameters is assumed. In the strict sense, prior distributions need to be inferred from an independent data source in a Bayesian framework, which is not always possible. A strong discrepancy between Bayesian and maximum-likelihood-based methods indicates that the underlying data set does not reflect the information from the prior distributions well, thereby identifying a potential problem.

Estimates of haplotype frequencies and MOI can be further utilized as plug-in estimates for population genetic inferences, such as detecting selection patterns or population structure. E.g., quantities such as heterozygosity or pattern of linkage-disequilibrium can be derived, which are informative on selection processes, such as in the case of drug resistance.

Standard population genetic analyses to mine patterns of selection are concerned with completed selective sweeps, i.e., the mutations of interest replaced the wildtypes. In the case of antimicrobial resistance, one is concerned with ongoing selective processes, i.e., not all variants of interest reached fixation in the population. Hence, signatures of selection are distorted. It is, therefore, necessary to condition population genetic analyses on the subpopulation which is characterized by variants of interest at certain markers (loci). Many methods to estimate MOI and variant frequencies are insufficient in this context, as they are applicable only in the case of too restrictive genetic architectures. This is true for methods that apply only to single markers or biallelic markers.

Here, the maximum likelihood methods from [10, 12] to estimate MOI and haplotype frequencies are extended to be applicable to a general genetic architecture. Because the method is capable of estimating frequencies of haplotypes characterized by multiple multiallelic loci, such estimates can be utilized for further population genetic analyses.

First, the underlying statistical model is derived under the assumption that MOI follows a conditional (positive) Poisson distribution (i.e., only disease-positive samples are considered). A number of illustrations help to facilitate the meaning of the mathematical notation. (Readers, with a limited mathematical background, can skip the formal details and move to the Results section, but are encouraged to be guided by the illustrations.) The model is complex to allow for a closed-form solution for the maximum likelihood estimate (MLE). Hence, the expectation-maximization (EM) algorithm is employed. The derivation of the algorithm is found in Deriving the EM-algorithm, and only the final iterative algorithm is presented in Results. Furthermore, expressions for haplotype prevalence are derived, which are epidemiologically more relevant than frequencies. Particularly, the interplay between the MOI distribution, frequencies, and prevalence becomes clear from the formulae. Importantly, prevalence cannot be directly observed from molecular data, since typically only unphased haplotype information is available. Consequently, it is in general ambiguous which haplotypes (variants) are present in an infection. In ad hoc approaches, sometimes only unambiguous information is considered (e.g., [6, 7]) to derive estimates for haplotype prevalence/frequencies. To facilitate the relations with such methods, also formulae for the prevalence of haplotypes in unambiguous observations are derived.

Furthermore, the asymptotic covariances between model parameters and their transform (mean MOI; haplotype prevalence) are derived. Two alternative approaches are discussed in Mathematical Appendix.

The finite sample properties of the estimator are explored by numerical simulations. These complement the results of [10, 12, 25], which in special cases indicate that the proposed method has little bias and that the estimator’s variance is close to the Cramér-Rao lower bound.

Special emphasis is given to data application. Specifically, a *P*.*falciparum* data set concerned with markers associated with sulfadoxine-pyrimethamine (SP) resistance collected in Yaoundé, Cameroon is analyzed. First, the frequencies and prevalence of resistance-associated haplotypes are calculated. In a second step, heterozygosity and pairwise LD are determined across a range of microsatellite markers flanking the mutations of interest, conditioned on resistance-associated haplotypes that surpass a minimum frequency threshold of 10%. The data application is intended to showcase possible applications of the proposed method and to discuss its limitations.

An implementation of the method as R script alongside detailed documentation is provided as supplements and is available via GitHub at https://github.com/Maths-against-Malaria/generalModel.git, where the material will be maintained, and Zenodo (DOI:) at [26].

## Results

First, the maximum likelihood estimates (MLE) of haplotype frequencies and the MOI parameter for the proposed statistical model are derived. Second, estimates of haplotype prevalences, which are epidemiologically and clinically more relevant quantities than frequencies, are derived using the MLEs as plugin estimates. Third, the asymptotic variance of the estimator is derived, i.e., the Cramér-Rao lower bound is derived as the inverse Fisher Information. This is achieved by two alternative approaches, shown to be equivalent: (i) by embedding the log-likelihood function in a higher dimensional space; (ii) by eliminating a redundant parameter. Next, the finite sample properties of the estimator are explored via numerical simulations. Finally, the use of the proposed method is demonstrated on a *P. falciparum* data set from Cameroon, to estimate heterozygosity and LD across a range of microsatellite markers on the background of haplotypes associated with SP-resistance.

### Maximum likelihood estimate

The estimates 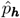 and 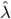 of haplotype frequencies and the MOI parameter are obtained by maximizing the log-likelihood function (53). Note that, even in the simple case of a single locus (in which the model can be written as an exponential family), no closed-form solution exists (cf. [9]), and one has to rely on numerical methods. Here, for this purpose, the expectation-maximization (EM)-algorithm is employed.

The EM-algorithm is an iterative procedure that alternates two steps: (i) the expectation (E) step during which the expectation, with regard to the parameter choice in the current step, of the log-likelihood function is calculated over an unobserved variable and the unknown model parameters; (ii) the maximization (M) step during which the function from the E step is maximized over the unknown parameters. This yields the parameter choice for the next E-step. The whole iteration is repeated until convergence, yielding the MLE. A detailed derivation of the algorithm is described in the Mathematical Appendix in section Deriving the EM-algorithm.

The algorithm begins with an initial choice for the MOI parameter 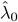 and haplotype frequencies 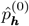. Given the parameter choices 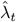 and 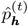 in step *t*, the estimates are obtained in step *t* + 1 by first updating the haplotype frequencies 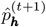 as

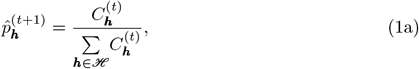

where

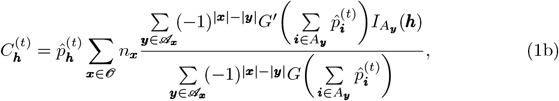

and

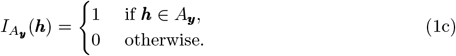

Then, the MOI parameter 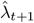 is obtained by iterating the following equation until convergence:

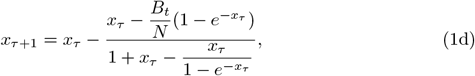

where

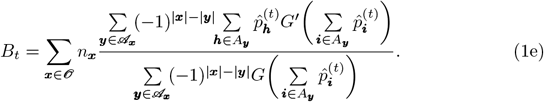

The iteration in (1d) starts with the initial value 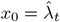 and is repeated until convergence, i.e., until |*x*_*τ*+1_ − *x*_*τ*_ | *< ε* is satisfied. The update of the Poisson parameter in step *t* + 1 is then 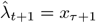.

The two iterative steps following (1) until numerical convergence, i.e., until 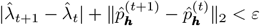. Once this holds, the MLE is obtained as

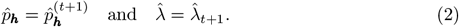

Note, the algorithm is guaranteed to converge to a point, at which a maximum of the original likelihood function is attained (not necessarily a global maximum), typically within a few iterations.

The mean MOI *ψ* is then estimated by evaluating (32b) at the MLE, i.e., as

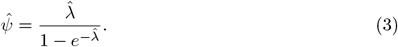

Considering the practical applicability, the difficulty here is to efficiently implement the rather complicated sums in (1). An implementation is available as an R script and comprehensive manual, provided as supplementary material, and a version subject to future updates can be accessed via GitHub at https://github.com/Maths-against-Malaria/generalModel.git, and Zenodo at [26].

### Estimation of prevalence

The prevalence of a haplotype is the probability that it occurs in an infection. Importantly, pathogen haplotypes are not directly observed in infections. Rather, the genetic information obtained from molecular assays is typically unphased, which leads to ambiguity if multiple haplotypes are present in an infection (cf. Fig. 10). Consequently, the absence and presence of certain haplotypes in an infection is uncertain.

Therefore, two concepts of prevalence are distinguished here: (i) “true prevalence”, which is the probability that a haplotype is present in an infection, and (ii) “conditional prevalence”, which is the probability that a haplotype occurs in an infection, in which it can be unambiguously observed. The distinction is important for practical purposes. In the first case, a statistical model, as the one proposed here, is required to resolve ambiguity in the observable information, while in the latter case, all samples with ambiguous information would be disregarded, and prevalence simply calculated as the relative frequency of the remaining samples, in which a particular haplotype occurs.

To distinguish between unobservable and conditional prevalence, it is necessary to emphasize that for an infection with MOI= *m*, two outcomes are possible: (i) a “single infection” involving the same pathogen haplotype *m* times, and (ii) a super-infection involving several pathogen haplotypes, i.e., “multiple infections”. Depending on the infecting pathogen haplotypes, multiple infections yield either ambiguous observations when the alleles of the infecting pathogen haplotypes are different at more than one marker locus or unambiguous observations if only the pathogen haplotype are infecting such that their alleles differ at exactly one marker locus. Note that single infections yield unambiguous observations.

In the following, we first derive the true prevalence, we then derive the conditional prevalence, which is conditioned on only observing unambiguous observations, i.e., single infections and unambiguous multiple infections.

#### True prevalence

For a given haplotype ***h***, let *q*_***h***_ denote its true prevalence. The derivations follow the steps in [12]. Moreover, assume an infection with observation ***x***, and the probability *q*_−***h***_ that haplotype ***h*** is not in ***x***. The true prevalence *q*_***h***_ is obtained as

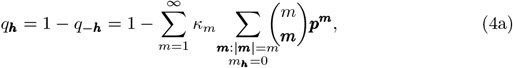

using the multinomial theorem, the innermost sum yields

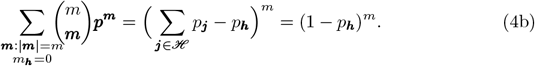

Hence,

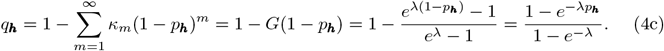

Therefore, given the MLEs 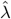 and 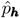 of the MOI parameter and haplotype frequencies, respectively, the true prevalence is estimated from (4c) by

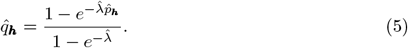

#### Conditional prevalence

In practice, it might be desirable to estimate haplotype prevalence based on observations in which haplotype information is unambiguous. In such a case, all ambiguous observations in the data set are discarded, and the prevalence of a given haplotype ***h*** is estimated as the probability of observing ***h***, conditioned on observing unambiguous observations.

The set of possible unambiguous observations is a subset of *𝒪* and is denoted by 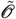, i.e., 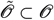. The conditional prevalence 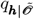 of haplotype ***h*** is defined as

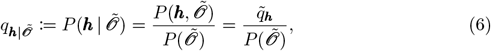

where, 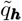 defines the probability that ***h*** is observed in an unambiguous observation, and 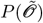 is the probability of all unambiguous observations.

For a given haplotype ***h***, to obtain the quantities 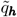 and 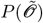, we need to construct the set *U*_***h***_ of all haplotypes ***i*** that yield unambiguous multiple infections with ***h*** (see Fig. 1). Recall that unambiguous multiple infections are obtained only for infections with exactly two pathogen haplotypes, such that their alleles are distinct at exactly one locus. Note that ***h*** is not in the set *U*_***h***_, i.e., ***h*** ∉ *U*_***h***_. Therefore, the set *U*_***h***_ is defined by

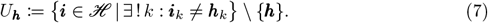

**Fig 1.**
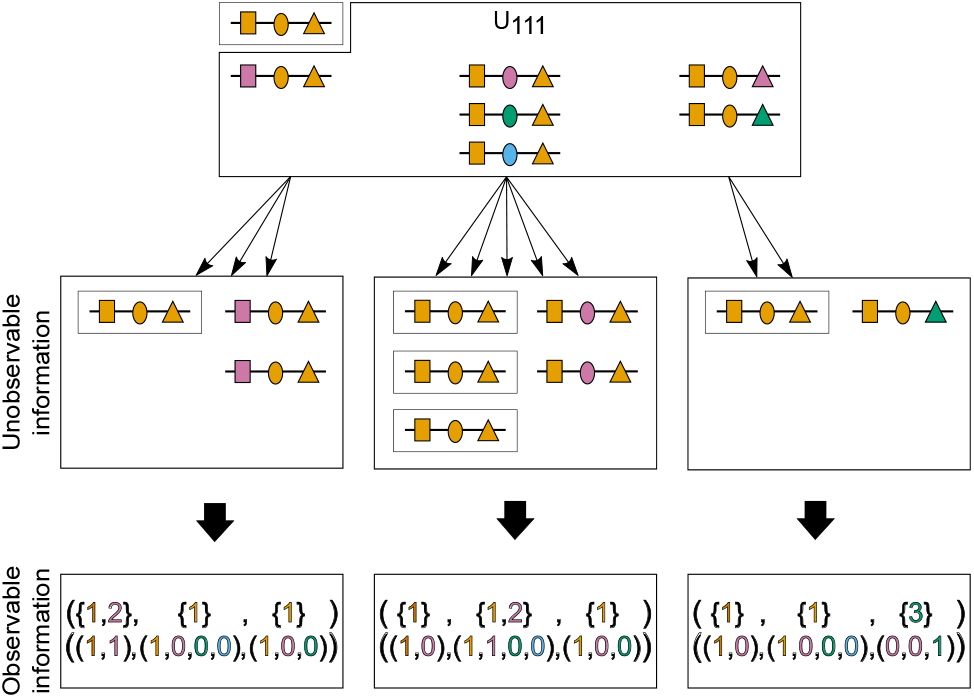
Illustration of unambiguity of infections between *h* and haplotypes in set *U*_*h*_: Shown are multiple infections involving haplotype ***h*** = (1, 1, 1), and (ii) a haplotype from the set *U*_111_ (upper block). In all resulting observations, haplotype information is unambiguous. However, MOI remains unknown.

The cardinality of *U*_***h***_ is 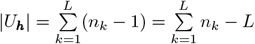, and for any haplotype ***i*** such that ***i*** ∈ *U*_***h***_, we have ***h*** ∈ *U*_***i***_.

The probability 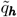 that haplotype ***h*** is observed in an unambiguous infection is the sum of the probabilities of all the multiple infections involving haplotype ***h*** and exactly one haplotype ***i*** in *U*_***h***_, and the single infection with ***h***. Assume a fixed haplotype ***h***, a haplotype ***i*** ∈ *U*_***h***_, and an unambiguous observation 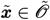 involving ***h***. For MOI= *m*, because ***i*** and ***h*** are sampled with replacement each of the *m* times, the following outcomes are possible for 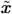: (i) ***h*** is sampled all *m* times, i.e., *m*_***h***_ = *m*, yielding a single infection with ***h***, (ii) ***i*** is sampled all *m* times, i.e., *m*_***i***_ = *m*, yielding a single infection with ***i*** and (iii) ***i*** and ***h*** are sampled *m*_***i***_ and *m*_***h***_ times, respectively, such that *m* = *m*_***h***_ + *m*_***i***_, yielding an unambiguous multiple infection (cf. [12]). Since unambiguous infections with ***h*** are of interest, only cases (i) and (iii) are relevant in the derivation of 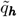. Therefore, one has

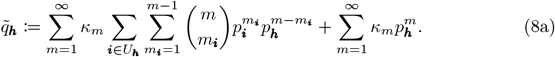

We show in section Prevalence estimates of the Mathematical Appendix that this quantity simplifies to

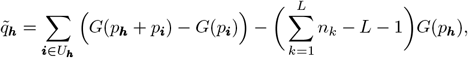

where *G* is the PGF (48b), and *L* is the number of loci considered.

The probability of all unambiguous observations 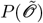 is derived in section Prevalence estimates of the Mathematical Appendix and is given by

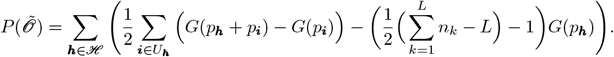

Hence, the conditional prevalence of ***h*** is given by

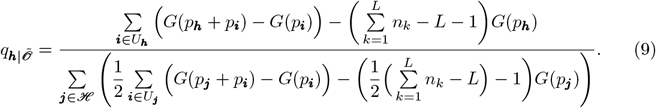

The estimate of conditional prevalence is obtained by plugging the MLEs in (9), i.e.,

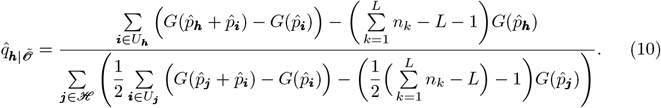

### Asymptotic variance and covariance

It is desirable to find unbiased estimators. In practice, this is hardly possible for complex statistical models. Specifically, maximum-likelihood estimators are typically only asymptotically unbiased, i.e., bias converges to zero as sample size increases to infinity. This also holds true for the present model, which was shown to be biased [10, 12, 22, 25]. For the special case of a single molecular marker, the method was shown to be asymptotically unbiased, because the model falls into the class of exponential families [22]. For the cases of two multiallelic markers and arbitrary many biallelic markers the model no longer falls into the class of exponential families and the estimator was shown numerically to be asymptotically unbiased [10, 12]. In any case, it is also desirable to derive estimators with small variance. For unbiased estimators, the minimum possible variance is given by the Cramér-Rao lower bound (CRLB) [27, 28]. Although biased estimators can have a lower variance, in practice, if an asymptotically unbiased estimator has variance close to the CRLB, there is not much hope to improve the estimator (cf. [29]). Hence, the CRLB is defined first, and it is shown numerically that it agrees well with the variance of the estimator. Together with the results of [10, 12, 22, 25] this provides a numerical proof that the MLE of the present model is efficient, i.e., its asymptotic variance reaches the CRLB.

#### Original model parameters

The CRLB is the inverse expected Fisher information [27, 28]. Given the model parameters ***θ*** and the log-likelihood function *L*_*𝒳*_ (***θ***) in (53), the Fisher information, denoted ℐ (***θ***), is the square matrix with entries

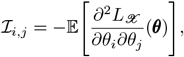

where *θ*_*i*_ and *θ*_*j*_ are components of the vector ***θ*** ∈ Θ = ℝ^+^ × *𝒮* _*H*_. The expectation is taken with regard to the distribution of the observations *P*_***x***_. In the context of this model, the parameter space is not an open set of ℝ ^*H*+1^, since the *H*-dimensional simplex is not an open set in ℝ ^*H*^, as one of the haplotype frequency, e.g., *p*_*H*_ can be written as a function of the others, i.e.,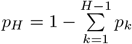. Results on the Fisher information require the parameter space to be an open set. Here, this can be resolved in two ways. First, one of the redundant parameters, e.g., *p*_*H*_, can be eliminated by substituting 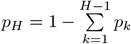.Second, the model can be embedded into a higher dimensional space by introducing a nuisance parameter.

Although the first approach seems simple at first sight, typically, calculations are simpler in the second approach. For the present model, this became evident in the special case of a single marker (cf. [9]). Hence, this approach is also pursued for the present general case, and it is formally proved that both approaches are equivalent (see section Fisher information matrix in the Mathematical Appendix in the supplementary material).

Following [9], we use a Lagrange multiplier as a nuisance parameter, by defining

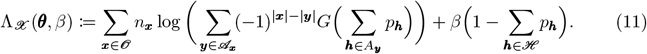

The Lagrange multiplier *β* guarantees that Λ_*𝒳*_ and the original likelihood function attain their maxima at the same point. Note, Λ_*𝒳*_ is not a log-likelihood function. It is defined formally for all points 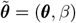 in an open real space, i.e.,

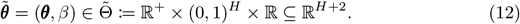

The equivalent of the Fisher information is defined as

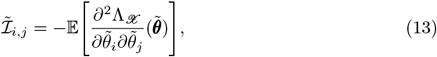

where *i* and *j* are the *i*-th and *j*-th element of the vector 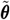. The expectation is formally taken with regard to *P*_***x***_, which in the higher-dimensional embedding might not be a probability distribution. Importantly, for inverting the above matrix, the rows and columns corresponding to the nuisance parameter *β* need to be considered.

A detailed description of the derivation of the entries of the Fisher information is presented in section Entries of information matrix in higher dimensional space in the Mathematical Appendix. The entries are as follows

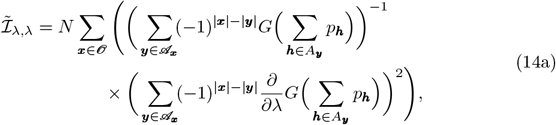

where

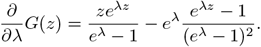

Additionally,

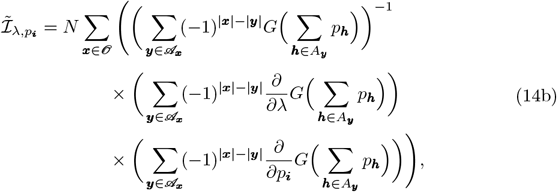

with

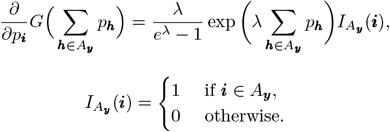

Moreover,

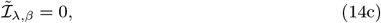

and

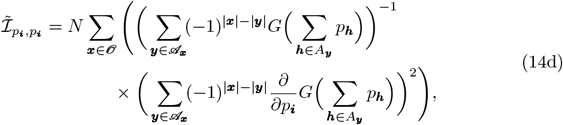

One also has

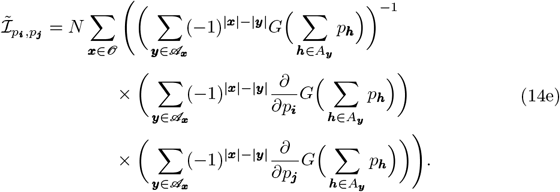

Finally,

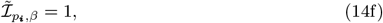

and

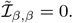

Hence, the “Fisher information matrix” has the form

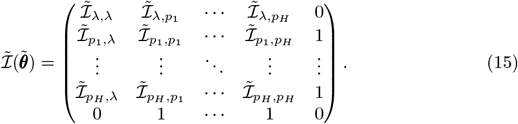

The results in Fisher information matrix in the Mathematical Appendix show that for any likelihood function, whose parameter space includes a simplex, the CRLB is given by the inverted high-dimensional embedded “Fisher information matrix” with the rows and columns corresponding to the nuisance parameters being disregarded after inversion. Here, this corresponds to disregarding the row and column corresponding to *β*. This is denoted by

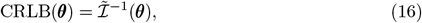

where the latter indicates that the row and column corresponding to *β* are disregarded, and the resulting matrix is evaluated at an admissible parameter ***θ*** ∈ Θ. This gives the asymptotic covariance matrix of the original model parameters *λ* and ***p***.

#### Mean MOI and haplotype frequencies

In practice, the mean MOI *ψ* is of more interest than the MOI parameter *λ*. The inverse Fisher information can be readily used to calculate the asymptotic covariance matrix of the mean MOI and haplotype frequencies. Namely, recall that 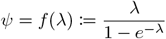. Let

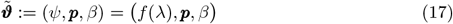

denote the parameters of interest. Thus, the log-likelihood function in terms of the new parameters becomes

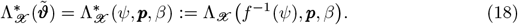

The “Fisher information” of the transformed parameter 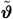 is defined as

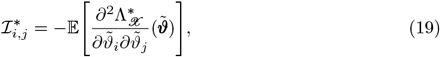

where *i* and *j* are the *i*-th and *j*-th element of 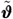. Straightforward application of the chain rule gives

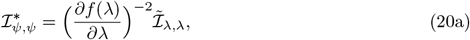

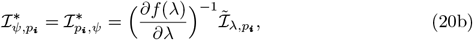

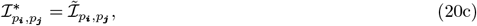

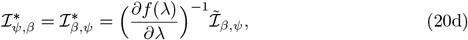

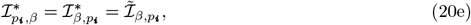

and

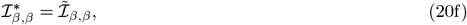

where

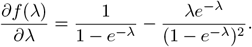

In compact form, this is written as

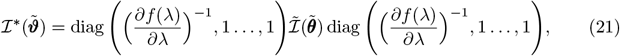

from which it is evident that the inverse matrix becomes

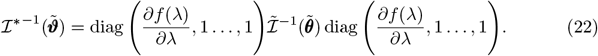

The asymptotic covariance matrix is again given by dropping the row and column corresponding to *β* in (22).

#### Mean MOI and prevalences

In practice, haplotype prevalences are sometimes of more interest than frequencies. The asymptotic variance of prevalences is derived from that of the original parameters by similar transformations as above. Namely, recall that the (true) prevalence of haplotype ***h*** is given by 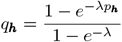. Let, mean MOI and prevalences can be expressed in terms of the original parameters as

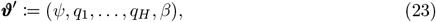

be the transformed parameters. These can be expressed in terms of the original parameters as

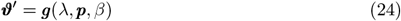

where the parameter transformation can be written as

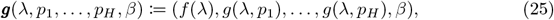

with *f* defined as in (32b) and

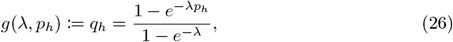

(recall the equivalence between haplotypes ***h*** and their rank *h*.)

The log-likelihood function expressed in terms of the transformed parameters becomes

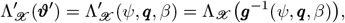

where ***q*** := (*q*_1_, …, *q*_*H*_).

As above, application of the chain rule yields the entries of the “Fisher information” of the transformed parameters. Particularly,

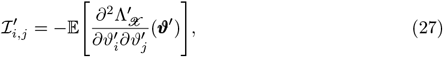

where *i* and *j* are the *i*-th and *j*-th element of the transformed parameter vector ***ϑ***^′^. By using the chain rule, one obtains

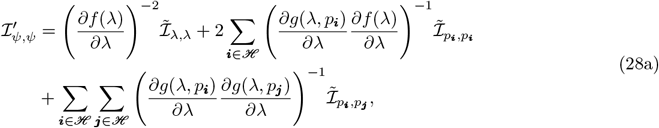

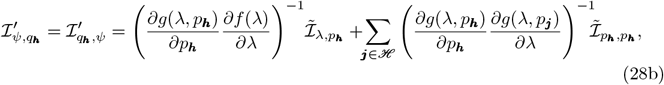

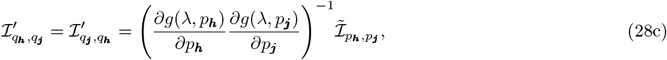

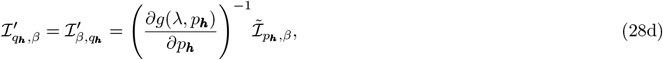

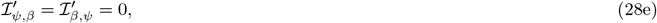

and

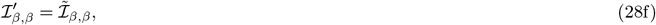

where

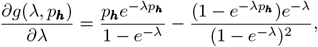

and

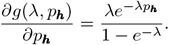

Note that the Jacobian matrix of the transformation ***g*** : ℝ^*H*+2^ → ℝ^*H*+2^, with 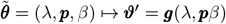, denoted by *J*, allows to rewrite ℐ^′^ as

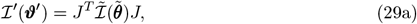

where

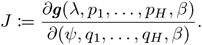

More explicitely,

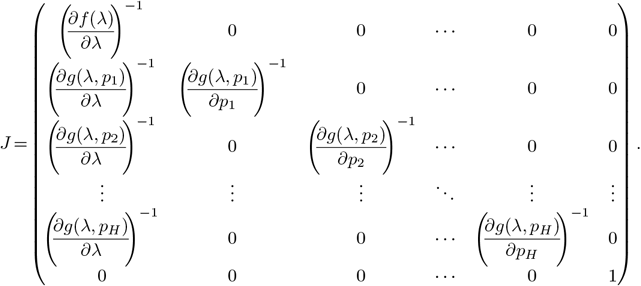

Therefore, the inverse of the matrix ℐ^′^ in (29a) is obtained as

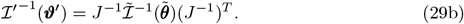

The inverse *J*^−1^ of the Jacobian matrix *J* can be obtained by blockwise inversion. First, rewrite *J* as

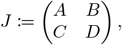

where

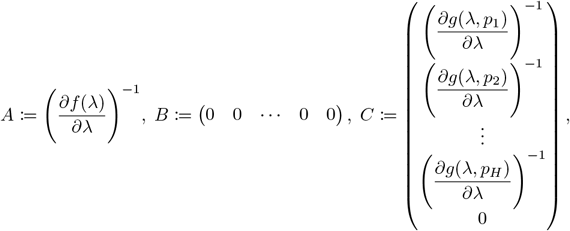

and

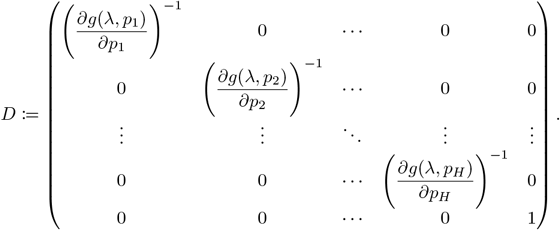

Blockwise inversion (cf. [30]) gives

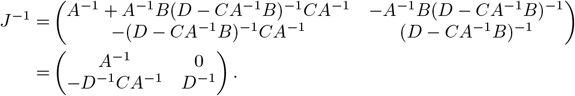

In the present case, this becomes

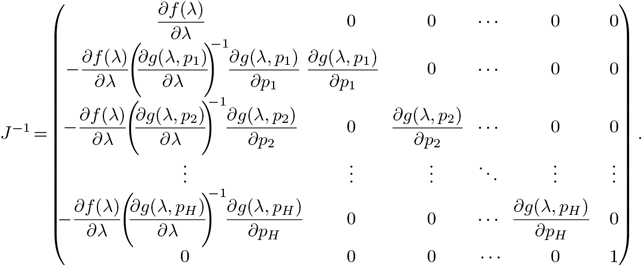

The inverse ℐ^′−1^ is the covariance matrix of the estimator ***ϑ***^′^ = ***g***(*λ, p, β*) and is obtained from (29b).

### Finite sample properties

Typically, maximum-likelihood estimators have convenient statistical properties. The proposed method was shown to have little bias under finite sample assuming the following genetic architectures: multiple biallelic loci [12], a single multiallelic locus [25], and two multiallelic loci [10]. Furthermore, for a single multiallelic locus the method was proven to be asymptotically unbiased as it falls within the class of exponential families [22]. In the general case of multiple multiallelic loci, due to the curse of dimensionality, it is impractical to systematically investigate bias, besides that it would add little to what is already known. Hence, here the finite sample properties of the variance of the estimator are investigated and compared with the asymptotic results of the last section (Asymptotic variance and covariance). Thus, the focus here is on the efficiency of the estimator. To make the discrepancy between the actual variance and the asymptotic predictions of the mean MOI comparable across the parameter range, variation is studied in terms of the coefficient of variation (CV) for mean MOI.

Numerical simulations yield a close agreement of estimated coefficient of variation of both the mean MOI and haplotype frequencies and their theoretical prediction (based on the CRLB) for sample size *N >* 50 (Figs. 2 and 3). For a small sample size of *N* = 50 a higher discrepancy is observed, particularly for more complex genetic architectures (compare Figs. 2A, C, and Figs. 2B, D). Namely, a larger number of alleles per locus yields an increased number of haplotypes, such that these are likely to have an inaccurate representation of the true haplotype distribution in a data set with a small sample size (*N* = 50). Consequently, the estimate is sensitive to outliers, resulting in high variance for the MOI estimate. Additionally, the variance of the haplotype frequency estimates increases (compared with the asymptotic approximation; see Fig. 3).

**Fig 2.**
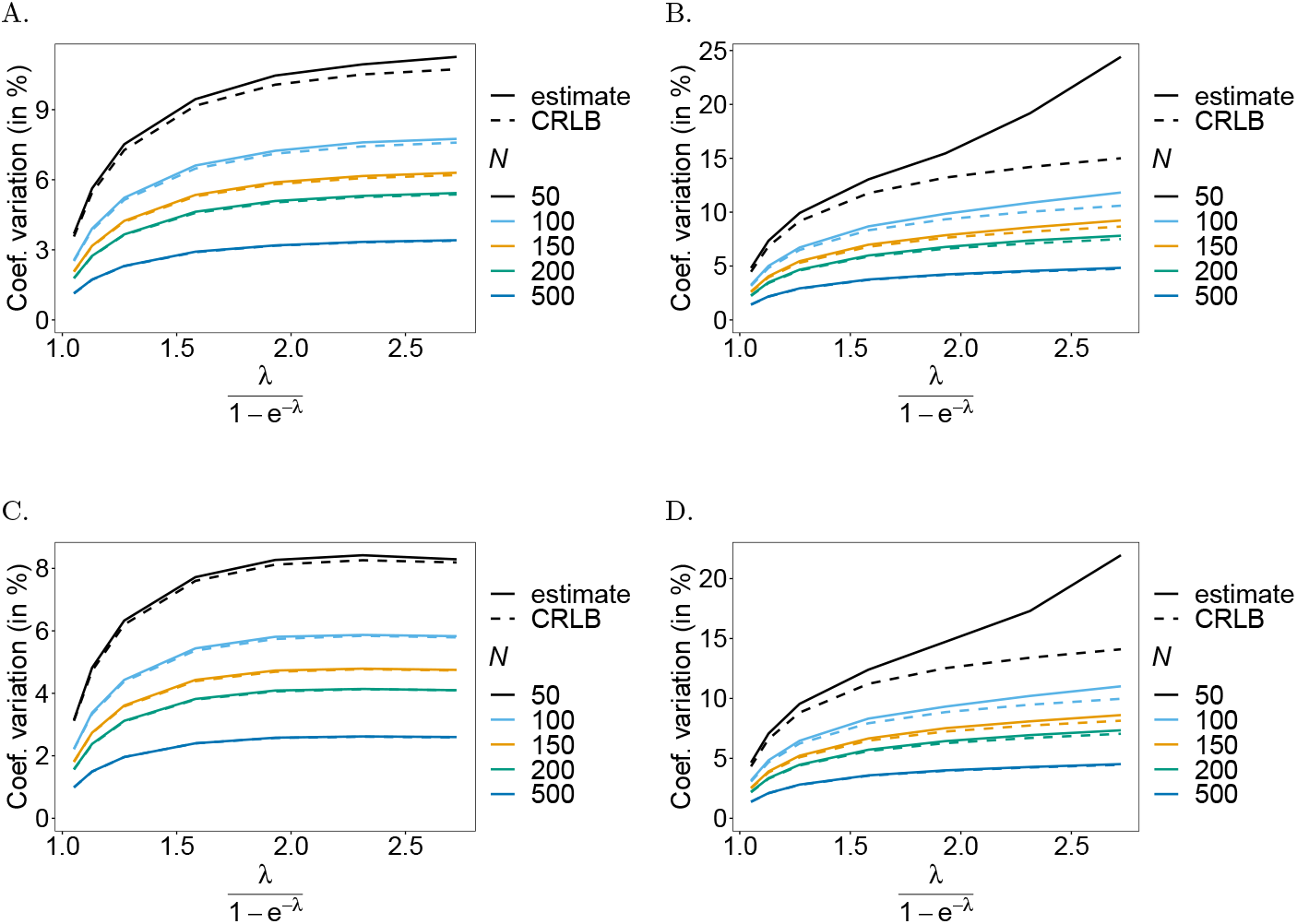
Cramér-Rao lower bound for MOI estimates: Shown is the coefficient of variation (CV) of the mean MOI estimates *ψ* in % as a function of the true mean MOI (i.e., for a range of MOI parameters). The genetic architecture is *n*_1_ = *n*_2_ = 2 in (A, C) and *n*_1_ = 4, *n*_2_ = 7 in (B, D). The frequency distribution in (A, B) is balanced while that in (C, D) is unbalanced. The different sample sizes are represented by the colors, the continuous lines are the CV from the MOI estimates and the dashed lines are the CV from the Cramér-Rao lower bound.

**Fig 3.**
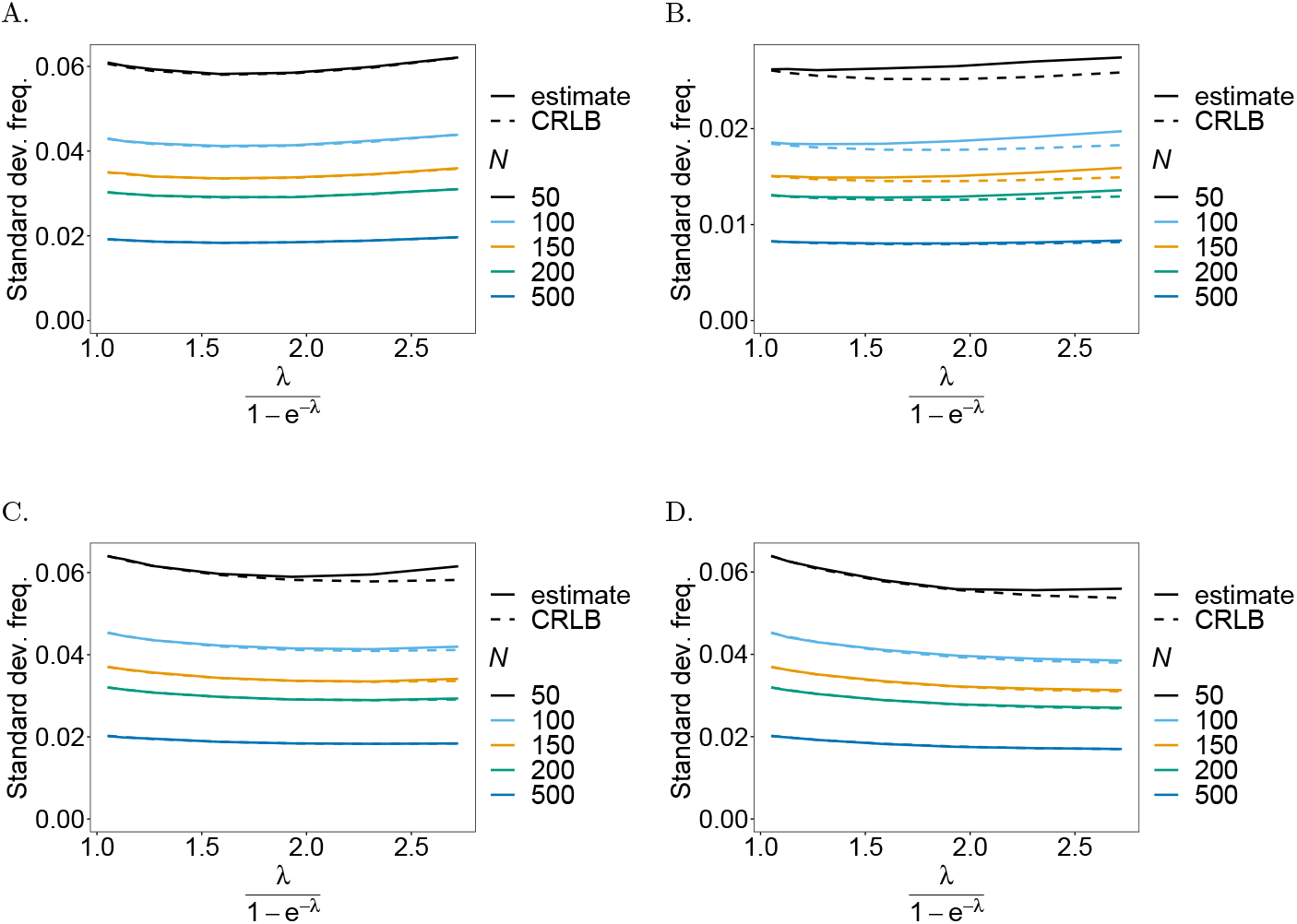
Cramér-Rao lower bound (CRLB) for haplotype frequencies estimates: Shown is the standard deviation (continuous line) alongside the CRLB (dashed line) for the estimates of haplotype frequencies as a function of the true mean MOI (i.e., for a range of MOI parameters). Panels (A, C) correspond to the estimation of a haplotype frequency whose true frequency is *p* = 0.25 assuming the genetic architecture *n*_1_ = *n*_2_ = 2. The case of the genetic architecture *n*_1_ = 4, *n*_2_ = 7 for a dominant haplotype with true frequency *p* = 0.7 is shown in panels (B, D). Notably, the true haplotype frequency distribution in (A, B) is balanced while that in (C, D) is unbalanced. The colors indicate the different sample sizes.

In summary, the numerical investigations suggest that the estimators of haplotype frequencies and MOI are efficient, and the asymptotic variance is properly predicted by the CRLB if sample size is sufficiently large (*N* ≥ 100). Note that in the context of malaria, sample sizes of *N* = 100 − 500 are realistic.

### Data application

Estimates of haplotype frequencies and MOI can be utilized for further population genetic analyses. This is illustrated using a *P. falciparum* data set from Yaoundé, Cameroon [31]. The data was collected in 2001, 2002, 2004, and 2005. In the following analysis, the data is stratified into two groups: years 2001-2002 (*N* = 166), and 2004-2005 (*N* = 165). The data set consists of markers associated with resistance against sulfadoxine-pyrimethamine (SP) in *P. falciparum*. Resistance to the fast-acting component pyrimethamine is determined by mutations at codons 51, 59, 108, and 164 in the *Pfdhfr* gene, while resistance to the slow-acting partner drug sulfadoxine is determined by mutations at codons 436, 437, 540, 581, and 613 in the *Pfdhps* gene [31].

The following analysis aims to estimate heterozygosity and pairwise-LD across 18 microsatellite markers on chromosomes 4 around *Pfdhfr*, 15 microsatellite markers around *Pfdhps*, as well as 8 neutral markers on chromosomes 2 and 3 [31]. Such analyses are useful to understand the spread of antimalarial drug resistance.

In standard population-genetic analyses, one considers past evolutionary processes, i.e., the variants under selection already reached fixation. This is different in the case of antimalarial drug resistance. Namely, one is interested in ongoing evolutionary processes, in which the mutations of interest did not yet reach fixation. Hence, all analyses need to be conditioned on the variants of interest. Otherwise, patterns of genetic hitchhiking and LD would be obstructed. In the present case, the variants of interest are haplotypes determined by *Pfdhfr* and *Pfdhps* mutations. To obtain a reasonable sample size, the following analyses are conditioned on haplotypes surpassing a threshold frequency of 10%.

To determine the frequencies of *Pfdhfr* and *Pfdhps* haplotypes, their frequencies were first estimated with the proposed method.

#### Frequency of resistance-associated haplotypes and MOI

The frequencies of *Pfdhfr* and *Pfdhps* haplotypes at the two-time points are given in Table 1. At *Pfdhfr*, only the triple-mutant 51**I**/59**R**/108**N**/I164, conferring strong resistance, was detected with a frequency higher than 10% at both time points. This suggests widespread resistance to SP in Yaoundé from 2001 to 2005. At *Pfdhps*, the haplotypes 436**A**/A437/K540/A581/A613, S436/437**G**/K540/A581/A613 were detected at a frequency higher than 10% in the years 2001-2002, while the double-mutant haplotype 436**A**/437**G**/K540/A581/A613 also surpassed the 10% threshold in 2004-2005 suggesting ongoing selection for sulfadoxine resistance.

**Table 1.** Frequencies estimates of *P. falciparum* haplotypes associated with SP-resistance from data collected in Cameroon in years 2001-2002, and 2004-2005. Haplotype frequencies above the 10% threshold are marked with an asterisk (^*^) in both year groups.

| Years | *Pfdhfr* |  |  |  |  |  | *Pfdhps* |  |  |  |  |  |  |
| --- | --- | --- | --- | --- | --- | --- | --- | --- | --- | --- | --- | --- | --- |
|  | Haplotype (codons) |  |  |  | Freq. (%) | 95% CI | Haplotype (codons) |  |  |  |  | Freq. in % | 95% CI |
|  | 51 | 59 | 108 | 164 |  |  | 436 | 437 | 540 | 581 | 613 |  |  |
| 2001-2002 | N | C | S | I | 9.7 | 5.9 – 13.7 | S | A | K | A | A | 6.2 | 3.0 – 9.9 |
|  | N | C | **N** | I | 2.0 | 0.4 – 4.0 | **A** | A | K | A | A | 33.6* | 27.4 – 40.3 |
|  | N | **R** | S | I | 0.4 | 0.0 – 1.3 | S | **G** | K | A | A | 45.9* | 38.9 – 52.9 |
|  | **I** | C | **N** | I | 2.1 | 0.4 – 4.1 | S | A | K | A | **S** | 2.0 | 0.3 – 5.0 |
|  | N | **R** | **N** | I | 9.8 | 6.2 – 13.7 | **A** | **G** | K | A | A | 7.5 | 4.0 – 11.4 |
|  | **I** | **R** | **N** | I | 76.0* | 69.9 – 81.9 | **A** | A | K | A | **S** | 1.3 | 0.0 – 2.9 |
|  |  |  |  |  |  |  | S | **G** | K | A | **S** | 2.7 | 0.4 – 5.1 |
|  |  |  |  |  |  |  | S | **G** | K | **G** | **S** | 0.4 | 0.0 – 1.3 |
| 2004-2005 | N | C | S | I | 2.6 | 0.7 – 4.9 | S | A | K | A | A | 6.3 | 3.1 – 10.0 |
|  | N | C | **N** | I | 0.9 | 0.0 – 2.4 | **A** | A | K | A | A | 27.6* | 21.6 – 33.5 |
|  | **I** | C | S | I | 0.9 | 0.0 – 2.3 | S | **G** | K | A | A | 50.4* | 43.2 – 57.6 |
|  | **I** | C | **N** | I | 0.4 | 0.0 – 1.4 | S | A | K | A | **S** | 0.8 | 0.0 – 2.4 |
|  | **I** | **R** | S | I | 2.6 | 0.9 – 4.5 | **A** | **G** | K | A | A | 14.5* | 9.4 – 19.9 |
|  | N | **R** | **N** | I | 3.9 | 1.6 – 6.9 | **A** | A | K | A | **S** | 0.4 | 0.0 – 1.9 |
|  | **I** | **R** | **N** | I | 88.7* | 84.0 – 93.0 |  |  |  |  |  |  |  |

The estimates of the MOI parameter and 95% bootstrap confidence intervals (CIs) are presented in Table 2. Estimates were obtained at both time points, based on *Pfdhfr* markers, and *Pfdhps* markers, as well as the combined *Pfdhfr* and *Pfdhps* markers. The merit of the latter is the inclusion of more information, at the cost of a deflated sample size due to missing data. Because the narrowest CIs are obtained in this case, the gain of information outbalances the reduction in sample size. (This is not a general principle but will depend on the particular data set.) The widest CIs are obtained for the estimates based only on *Pfdhfr* markers. The reason is the unbalanced haplotype frequency distribution (cf. Table 1).

**Table 2.** MOI estimates from data collected in Cameroon in years 2001-2002, and 2004-2005.

| Years | *Pfdhfr* |  | *Pfdhps* |  | *Pfdhfr* & *Pfdhps* |  |
| --- | --- | --- | --- | --- | --- | --- |
|  | MOI | 95% CI | MOI | 95% CI | MOI | 95% CI |
| 2001-2002 | 0.991 | 0.696 – 1.361 | 0.772 | 0.566 – 0.999 | 0.94 | 0.749 – 1.152 |
| 2004-2005 | 0.752 | 0.407 – 1.267 | 0.799 | 0.580 – 1.055 | 0.865 | 0.651 – 1.099 |

Between 2001-2002 and 2004-2005, a slight reduction in MOI seems to be observed, although the CIs overlap by a great extent. Only for the estimates based on *Pfdhps* markers, there seems to be a very slight increase in MOI. This is presumably because the haplotype frequency distribution was less balanced in 2004-2005 (which also resulted in slightly wider CIs).

Note that it is preferable to estimate MOI from neutral markers rather than those under selection (cf. [25]).

#### Prevalence of resistance-associated haplotypes

For completeness, also estimates of haplotype prevalence are presented (Table 3). Due to the moderate value of the MOI parameters, frequency and prevalence are similar. Most discrepancies occur for the *Pfdhps* haplotypes in 2001-2002. This is clear from (4c) because the *Pfdhps* have the most balanced frequency distribution.

**Table 3.** Prevalence estimates of *P. falciparum* haplotypes associated with SP-resistance from data collected in Cameroon in years 2001-2002, and 2004-2005.

| Years | *Pfdhfr* |  |  |  |  |  | *Pfdhps* |  |  |  |  |  |  |
| --- | --- | --- | --- | --- | --- | --- | --- | --- | --- | --- | --- | --- | --- |
|  | Haplotype (codons) |  |  |  | Prev. (%) | 95% CI | Haplotype (codons) |  |  |  |  | Prev. in % | 95% CI |
|  | 51 | 59 | 108 | 164 |  |  | 436 | 437 | 540 | 581 | 613 |  |  |
| 2001-2002 | N | C | S | I | 14.5 | 9.1 – 20.0 | S | A | K | A | A | 8.7 | 4.3 – 13.8 |
|  | N | C | N | I | 3.1 | 0.6 – 6.1 | **A** | A | K | A | A | 42.5 | 34.9 – 50.1 |
|  | N | **R** | S | I | 0.6 | 0.0 – 2.0 | S | **G** | K | A | A | 55.5 | 47.7 – 63.0 |
|  | **I** | C | N | I | 3.3 | 0.7 – 6.4 | S | A | K | A | **S** | 2.8 | 0.5 – 7.1 |
|  | N | **R** | N | I | 14.7 | 9.5 – 20.5 | **A** | **G** | K | A | A | 10.4 | 5.6 – 15.7 |
|  | **I** | **R** | N | I | 84.2 | 78.5 – 89.6 | **A** | A | K | A | **S** | 1.8 | 0.0 – 4.2 |
|  |  |  |  |  |  |  | S | **G** | K | A | **S** | 3.8 | 0.6 – 7.2 |
|  |  |  |  |  |  |  | S | **G** | K | **G** | **S** | 0.6 | 0.0 – 1.8 |
| 2004-2005 | N | C | S | I | 3.6 | 1.1 – 6.6 | S | A | K | A | A | 9.0 | 4.5 – 14.3 |
|  | N | C | N | I | 1.2 | 0.0 – 3.1 | **A** | A | K | A | A | 35.9 | 28.6 – 43.6 |
|  | **I** | C | S | I | 1.2 | 0.0 – 3.1 | S | **G** | K | A | A | 60.2 | 52.5 – 68.1 |
|  | **I** | C | N | I | 0.6 | 0.0 – 1.9 | S | A | K | A | **S** | 1.2 | 0.0 – 3.5 |
|  | **I** | **R** | S | I | 3.7 | 1.2 – 6.8 | **A** | **G** | K | A | A | 19.8 | 13.5 – 26.7 |
|  | N | **R** | N | I | 5.5 | 2.4 – 9.2 | **A** | A | K | A | **S** | 0.6 | 0.0 – 2.6 |
|  | **I** | **R** | N | I | 92.1 | 87.8 – 95.8 |  |  |  |  |  |  |  |

Although the frequency of the single *Pfdhps* mutant haplotype S436/437**G**/K540/A581/A613 is below 50% in 2001-2002, its prevalence is estimated to be larger than 50%, implying that this variant was present in more than half of the infections.

#### Genetic hitchhiking

Conditioned on the SP-resistant *Pfdhfr* triple-mutant haplotype 51**I**/59**R**/108**N**/I164, heterozygosity estimates are shown for the years 2001-2002 and 2004-2005 in Figs. 4A and B, respectively. Heterozygosity was calculated for each microsatellite marker separately to retain the maximum possible sample size. (This is preferable because considering all microsatellite markers at the same time to calculate haplotype frequencies and then marginalizing to obtain allele frequency spectra for each marker would substantially deplete sample size. Namely, only samples without missing data at all loci could be retained. Nevertheless, analyzing markers separately still requires a method that allows for a general genetic architecture, since it is performed conditioned on *Pfdhfr* and *Pfdhps* haplotypes.)

**Fig 4.**
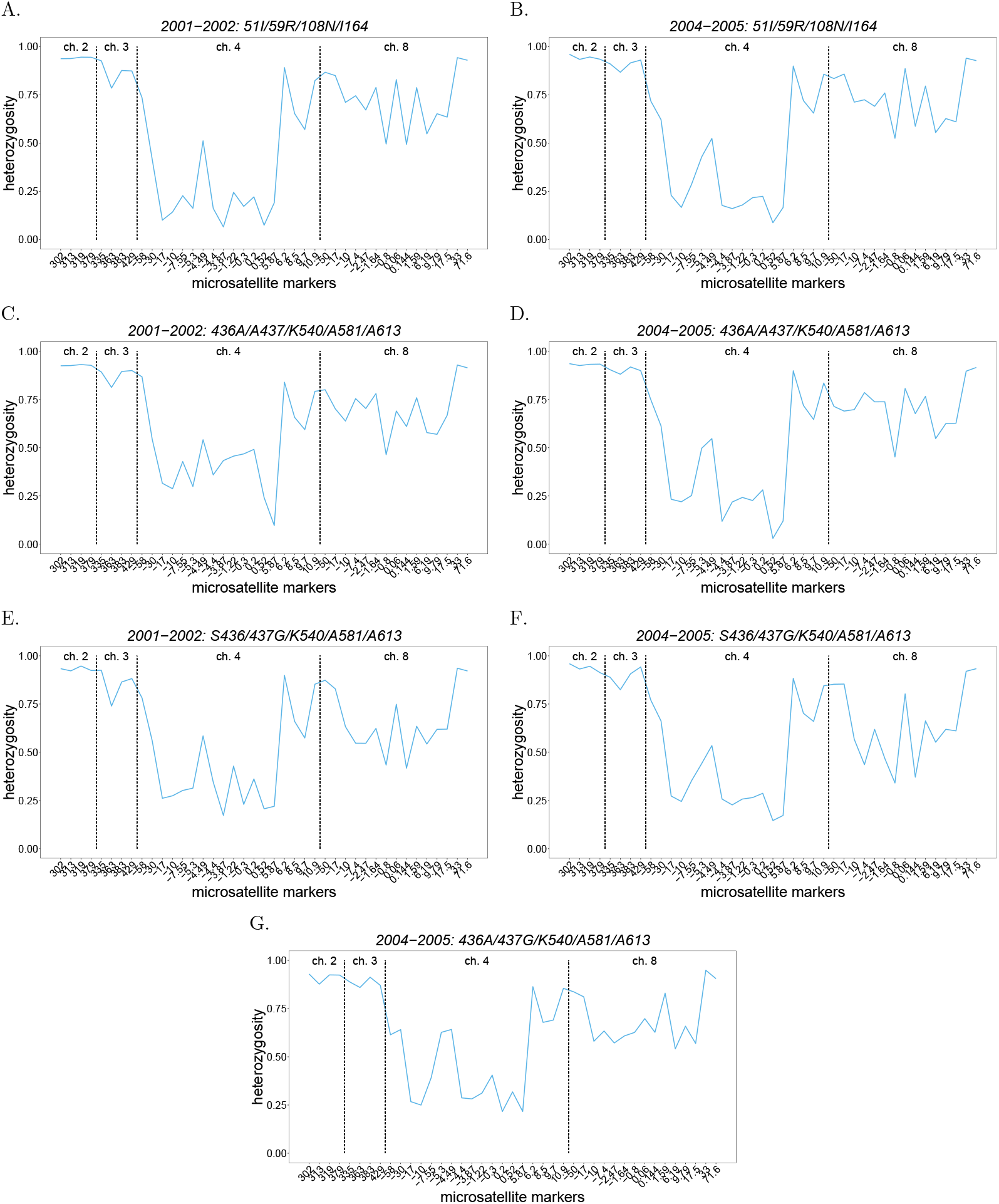
Conditional heterozygosity at microsatellite markers: Shown are estimates of heterozygosity across several microsatellite markers located on chromosomes 2 and 3, as well as on chromosome 4 surrounding *Pfdhfr* and chromosome 8 surrounding *Pfdhps* (chromosomes are separated by the vertical dashed lines) conditioned on various *Pfdhfr* and *Pfdhps* haplotypes either in the years 2001-2002 or 2004-2005 (shown at the top of each panel). Markers are ordered by their position in kilobases on the chromosomes. On chromosomes 4 and 8 the positions are relative to *Pfdhfr* and *Pfdhps*.

Heterozygosity at markers surrounding *Pfdhfr* is low, compared with that at more distant markers, exposing a clear selective sweep, i.e., traces of selection acting on *Pfdhfr*. Heterozygosity appears slightly higher in 2004-2005 (compare Figs. 4A and B). Although the differences are within the margin of error, heterozygosity is expected to increase over time. A drop in selection pressure could amplify this since treatment policy abandoned SP as first-line treatment in Cameroon in 2004 [31].

Such patterns are not as pronounced in *Pfdhps* but are still visible. The depletion in heterozygosity surrounding *Pfdhps* is slightly more pronounced around the S436/437**G**/K540/A581/A613 compared with the 436**A**/A437/K540/A581/A613 mutation at both time points, which corresponds to stronger selection for the S436/437**G**/K540/A581/A613 haplotype. The hitchhiking pattern flanking *Pfdhps* haplotypes corresponds to “soft selective sweeps” from recurrent mutations [32], i.e., the resistance-associated mutations emerged *de novo* or were imported several times at the background of different haplotypes. Such a soft-sweep pattern (with even higher heterozygosity) is also observed on the background of the 436**A**/437**G**/K540/A581/A613 double mutant (see Fig. 4G). It is plausible that the A → **G** mutation at codon 437 occurred on the background of the single-mutant 436**A**/A437/K540/A581/A613, while at the same time, the S → **A** mutation at codon 436 occurred on the background of the S436/437**G**/K540/A581/A613 haplotype, leading to soft-sweeps from independent origins, which therefore retains the levels of genetic variation.

The patterns of genetic hitchhiking in combination with the frequency distribution of *Pfdhfr* and *Pfdhps* haplotypes suggest stronger selection for pyrimethamine resistance. This is intuitive since pyrimethamine is considered the fast-acting component of SP and targeting *Pfdhps* with sulfonamides alone, would not be sufficient for malaria chemotherapy. Pyrimethamine was used as a monotherapy in the 1950s in some countries [33]. This suggests that resistance to pyrimethamine started to evolve earlier than resistance against sulfadoxine.

#### Pairwise linkage-disequilibrium

Conditional pairwise LD-values are obtained using *r*^2^ (and *D*^′^ as a comparison). LD is calculated at both time points for each pair of markers conditioned on the *Pfdhfr* triple mutant 51**I**/59**R**/108**N**/I164 at *Pfdhfr* (Figs. 5A, B), and the *Pfdhps* single mutants 436**A**/A437/K540/A581/A613 (Figs. 5C, D) and S436/437**G**/K540/A581/A613 (Figs. 5E, F), as well as on the *Pfdhps* double mutant 436**A**/437**G**/K540/A581/A613 at the second time point (Figs. 5G).

**Fig 5.**
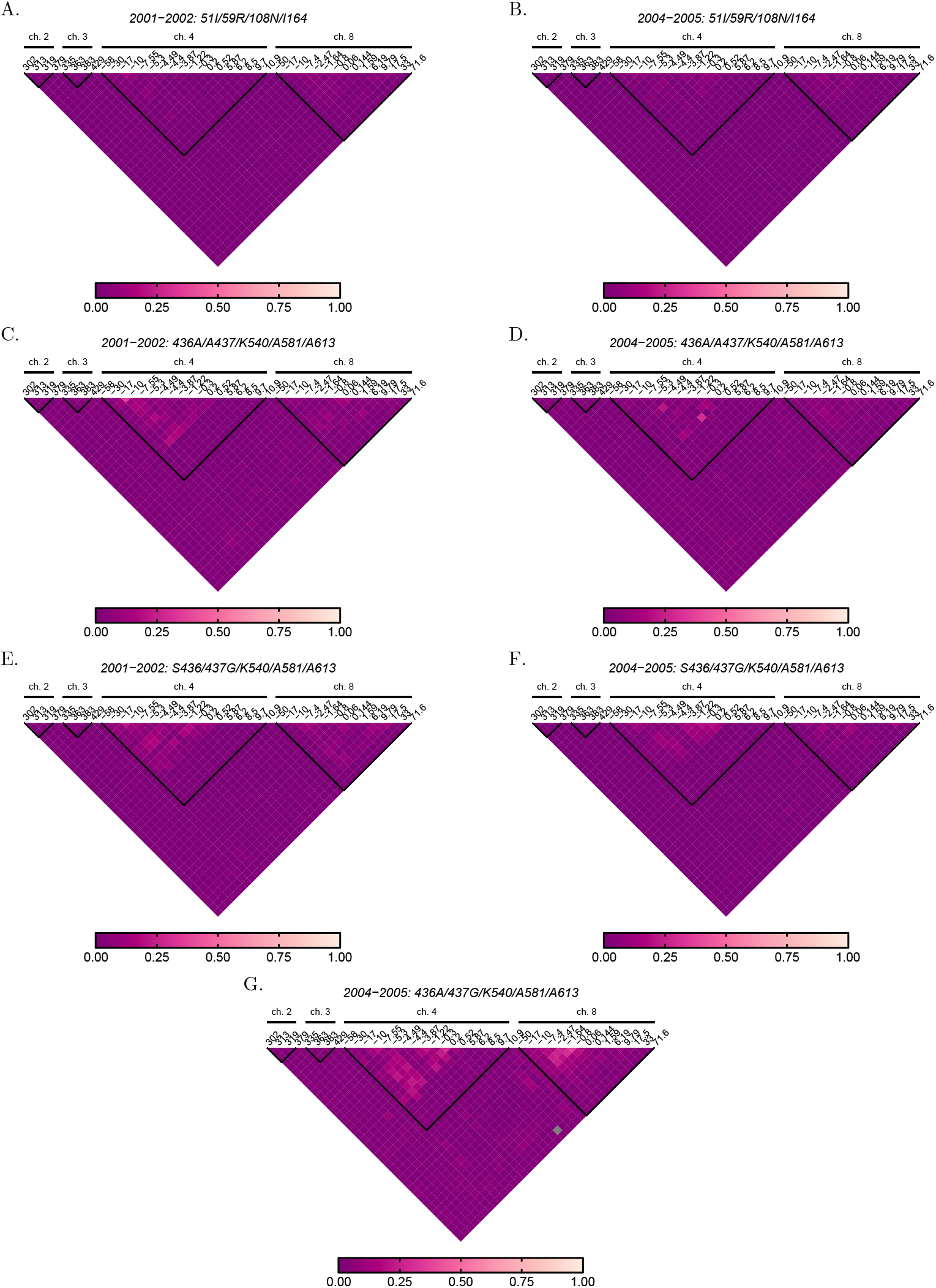
Conditional LD at microsatellite markers: As in Fig 4 but for pairwise-LD values calculated with *r*^2^.

Multiallelic LD measures are notoriously difficult to interpret and often uninformative. Conditioned on the *Pfdhfr* triple mutant, only slight LD is observed flanking *Pfdhfr* and *Pfdhps* (Figs. 5A, B). This suggests that the triple mutant was predominant for long enough to allow recombination to restore linkage equilibrium (LE).

Note that microsatellite markers are highly variable and fast-evolving, hence recombination and mutation will tend to restore genetic variation. This is particularly true given the results on heterozygosity, which suggest soft selective sweeps around *Pfdhps* variants. This is confirmed by looking at LD conditioned at the *Pfdhps* mutations. Conditioned on these, pairwise LD is more pronounced around *Pfdhfr* than around *Pfdhps*, on the one hand reflecting the soft sweep nature of *Pfdhps* mutations, on the other hand exposing signatures of selection around *Pfdhfr* (most haplotypes are linked to the triple mutant, which is predominant). This suggests that different variants flanking *Pfdhfr* were affected by the soft sweeps at *Pfdhps*. These patterns will overlap and camouflage the traces of selection when conditioning only on the *Pfdhfr* triple mutant. (Note that conditioning on joint *Pfdhfr* and *Pfdhps* haplotypes are impractical here due to missing data, which would substantially reduce sample size.)

The patterns of LD flanking *Pfdhfr* and *Pfdhps* are most pronounced conditioned on the *Pfdhps* double mutant, which (as it is mainly linked to the predominant *Pfdhfr* triple mutant) experiences the highest selective pressure.

Although multialellic LD in terms of *r*^2^ is hardly informative on the underlying evolutionary process in this case, it is at least intuitive. This is different for the *D*^′^ measure (Fig. 6). In fact, *D*^′^ leads to counterintuitive results, with the highest levels of LD between neutral markers at chromosomes 2 and 3, and lower LD flanking the loci under selection. These observations are purely an artifact of a small sample size. Namely, a large number of alleles are segregating at these markers. For example, two markers with, e.g., *n*_1_ = 20 and *n*_2_ = 25 alleles, would lead to 500 possible haplotypes. If all haplotypes had a frequency of 0.002, the markers would be in perfect LE. However, when taking a sample of size *N* = 165, the vast majority of haplotypes will not be sampled, while it is highly likely to sample all alleles at both markers. This leads to high LD values (i.e., most of the terms 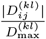 in (64a) equal to 1). Thus, *D′* will tend to overestimate LD for highly polymorphic markers unless sample size is unrealistically large. This problem does not occur around markers flanking targets of selection, because polymorphism is reduced, i.e., the number of alleles segregating at these markers is substantially lower. This artifact is also seen when comparing *D*^′^ conditioned on the *Pfdhfr* triple mutant *vs*. other mutants which are subject to weaker selection (Figs. 5A, B *vs*. C-G). Namely, genetic hitchhiking, which in the case of malaria can be genome wide [34], reduces polymorphism at neutral markers. This effect is more pronounced conditioned on variants under stronger selection, thereby mitigating the undesired behavior of *D*^′^ for highly polymorphic markers in small samples.

**Fig 6.**
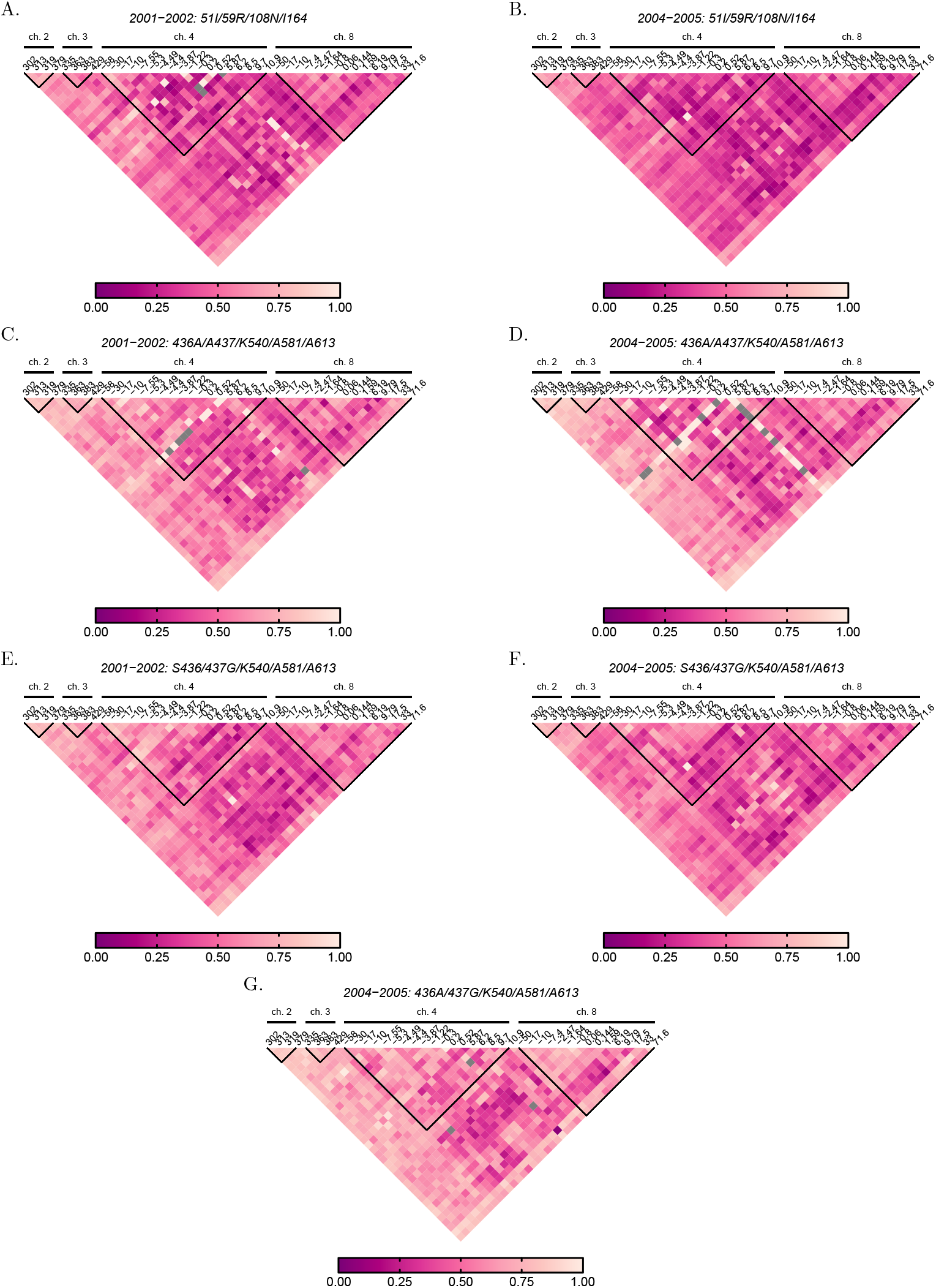
Conditional LD at microsatellite markers: As in Fig 4 but for pairwise-LD values calculated with *D*^′^.

## Discussion

Due to advancements in molecular/genetic assays over the past decades, molecular disease surveillance has become increasingly feasible and popular. Here, a method to estimate pathogen variants (haplotypes) frequencies and multiplicity of infection (MOI) was introduced.

The method extends those of [8, 10, 12] to accommodate a general genetic architecture, i.e., pathogen variants are characterized by an arbitrary number of multiallelic markers (loci). Following [8–10, 12, 35], MOI is defined as the number of super-infections with the same or different pathogen variants (cf. [2] for a detailed discussion). This only approximates co-infections, i.e., the co-transmission of different pathogen variants during the same infective event.

First, the underlying probabilistic model was derived. Because it does not allow a closed-form solution, the EM algorithm was employed to derive an efficient numerical iteration to derive the maximum likelihood estimates. Furthermore, formulae for prevalence were derived.

Whereas frequency refers to the relative abundance of a pathogen variant in the pathogen population, prevalence refers to the probability that the variant is present in an infection. Prevalence is more relevant for clinical and epidemiological considerations. Although the terms prevalence and frequency are sometimes used synonymously in the literature, they are different. In the absence of multiple infections, frequency and prevalence coincide. However, due to MOI prevalence always exceeds frequency (cf. [2]). Commonly, molecular/genetic data is not phased, i.e., haplotype information is missing. As a consequence, molecular data is ambiguous regarding the haplotypes actually present in an infection. In the case of malaria, often heuristic ad-hoc approaches are used to derive haplotype prevalence based only on unambiguous information. To relate the formal statistical approach to these heuristic ones, formulae for prevalence from unambiguous observations were derived.

The asymptotic covariances, i.e., the inverse Fisher information, were derived for the model parameters (haplotype frequency and MOI parameter) and their transforms in terms of prevalence and mean MOI. Although the Fisher information was derived explicitly, it was not possible to provide analytic expressions for its inverse. Importantly, two alternative approaches to derive the Fisher information were employed. First, a nuisance parameter was introduced to embed the parameter space into a higher dimensional open set in a real space. Second, a redundant parameter was eliminated (the haplotype frequencies are elements of the simplex and hence dependent), to reduce the parameter space to an open set in a lower-dimensional real space. This leads to different information matrices. It was shown that both approaches are equivalent and that corresponding entries of the inverse information matrices coincide.

By numerical simulations, [12] and [10] already showed that the method has little bias in special cases, which vanishes with increasing sample size, suggesting that the method is asymptotically unbiased. Here, these results were complemented, focusing on the bias of haplotype frequencies. Furthermore, it was numerically verified that the covariances of the estimator are well approximated by the inverse Fisher information (Cramér-Rao lower bound; CRLB). Simulations showed that the variance of the estimator (for haplotype frequencies and MOI) agrees well with the CRLB even for moderate sample sizes (*N*≥ 100). The largest discrepancies occur if sample size is small (*N <* 100) and MOI is high. This is less problematic in practice since larger sample sizes should be feasible in high transmission areas. The CRLB defines the minimum variance of an unbiased estimator. The fact that the covariance matrix of the estimator is well approximated by the inverse Fisher information for finite sample size, suggests that the proposed method has desirable asymptotic properties.

Application of the method was illustrated by analyzing a *Plasmodium falciparum* data set consisting of genetic markers associated with resistance to sulfadoxine-pyrimethamine (SP), caused by point mutations in the *Pfdhfr* and *Pfdhps* genes. First, the frequencies and prevalence of resistance-associated haplotypes were derived. In a second step, heterozygosity and pairwise linkage disequilibrium (LD) across microsatellite markers flanking the loci associated with resistance were calculated. These calculations were performed conditioned on specific *Pfdhfr* and *Pfdhps* haplotypes. These analyses were performed to showcase that the MOI and haplotype frequency estimates can be utilized for further population genetic analysis. The example also highlighted the pathological behavior of one multiallelic measure of LD (*D*’).

Importantly, the analysis required estimates of haplotypes, determined by a general genetic architecture (multiple mutiallelic markers). The special cases of the proposed method published earlier [9, 10, 12] are insufficient for such analyses.

The method is designed for haploid pathogens. This is hardly a restriction given that many pathogens, including viruses, bacteria, protozoa, etc., are haploid. Nevertheless, the method can be extended to be applicable to diploid or polyploid pathogens, although the relevance of such generalizations is limited.

Besides the promising properties of the proposed method, there are several shortcomings. Unlike in the case of a single molecular marker, there is no formal proof of existence, uniqueness, asymptotic unbiasedness, consistency, and efficiency. For a genetic architecture of a single molecular marker, the model can be rewritten as a natural steep exponential family, which proves the desired properties (importantly existence and uniqueness holds except in degenerate cases, e.g., if one variant is found in all samples). For two or more markers, the sum in (48a) does not factorize accordingly. However, the agreement between the CRLB and the variance of the estimator determined by simulations suggests that the estimator is efficient. Moreover, the fact that bias decreases with sample size, suggests asymptotic unbiasedness (which also suggests uniqueness of the estimator).

MOI is assumed to follow a conditional Poisson distribution, which might be inappropriate if MOI is over-dispersed. Intuitively, a negative binomial distribution seems more appropriate. However, in [19] it was pointed out that the likelihood estimation of negative binomial data is problematic, as the model seems to have degenerate behavior. (This readily occurs for maximum likelihood estimation of negative binomially distributed data. Specifically, the MLE will yield the limit of a Poisson distribution, if the data is not sufficiently over-dispersed.) As an alternative, [19] assumed a non-parametric distribution of MOI, which is the most flexible model. Except, for extreme cases this model did not outperform the Poisson model, suggesting that the assumption of Poisson-distributed MOI is justified.

The estimators can be improved by applying bias correction as in [21] in the case of a single marker. This led to an improvement in the bias of the MOI parameter (the gain for marker frequencies is marginal, as these are hardly biased). Such bias corrections might be tedious in the present model. As an alternative, parametric or non-parametric bootstrap bias corrections can be applied (cf. [36]).

Another shortcoming of the model is its inability to handle missing data. Namely, observations that lack information at one or more markers need to be disregarded. This yields a tradeoff between the confidence in the estimates and the number of molecular markers included. Namely, while the addition of more markers increases information, it also deflates sample size due to missing data and increases the number of model parameters geometrically. For a single marker, [22] proposed a model that incorporates missing information. This method was only superior to ignoring missing information if these were common. Incorporating missing information for a general genetic architecture leads to combinatorial difficulties in practice. In particular, all possible observations would need to be considered. Assuming 10 markers with 10 alleles per marker yields 1.25 ×10^3^0 possible observations. For computational feasibility, restrictions, and approximations would need to be applied to the probabilistic model to render it numerically feasible. An alternative to incorporate missing data would be to impute missing information. This can be readily achieved by estimating the model parameters first by disregarding missing information, and then use these estimates to impute missing information for each sample separately based on the marginalization of the distribution over the markers with missing information. In the final step, the estimates would be updated by those of the imputed data set.

Concerning numerical feasibility, the method is appropriate for a moderate number of molecular markers, with a moderate number of alleles per marker. Whether the method is feasible for a particular genetic architecture depends crucially on the underlying data set. As an example, assume 20 biallelic SNPs. Further assume, that only two haplotypes are present, the wildtype and mutant (these differ at all 20 SNPs). If both haplotypes occur in an infection, the formula for the resulting observation considers all 1048576 possible haplotypes, which give rise to more than 3.5 billion summands in equation (48a). In practice, such observations do not occur, and a genetic architecture of more than 20 SNPs is feasible.

In any case, there are restrictions to the generic architecture in practice. Particularly, the supported genetic architectures are insufficient to study relatedness of haplotypes within an infection (cf. [37–42]). Relatedness is important to distinguish super-from co-infections. Here, co-infections are ignored and only approximated by super-infections (cf. [25] for a more detailed discussion).

Here, a maximum-likelihood approach was pursued. Bayesian alternatives can be readily implemented and should be in good agreement as long as an uninformative prior distribution is used.

Besides the limitations, the proposed method is valuable for population genetic analysis of infectious diseases similar to malaria for a limited amount of molecular data, which is frequently collected in practice. An efficient implementation of the model is provided as an R script in the supplement, on GitHub https://github.com/Maths-against-Malaria/generalModel.git and Zenodo at https://doi.org/10.5281/zenodo.14452432 (cf. [26]). Moreover, detailed documentation of the R script with examples is provided.

## Methods

### Model background

The objective is to estimate the distributions of multiplicity of infection (MOI) and pathogen lineages for infectious diseases similar to malaria. Here, maximum-likelihood estimation is used. Although the following is not confined to malaria, this example is used to motivate the statistical model.

Here, the term multiplicity of infection refers to the number of super-infections during one disease episode, i.e., to the number of independent infectious events with potentially different variants of the same pathogen (see Fig. 7A). Importantly, it is assumed that during an infectious event, only one pathogen variant is transmitted. This is different from co-infections which refers to the co-transmission of several pathogen variants during one infectious event (see Fig. 7B). Importantly, super-infections with different pathogen variants rather than with different pathogens are considered (cf. Figs. 7A with C).

**Fig 7.**
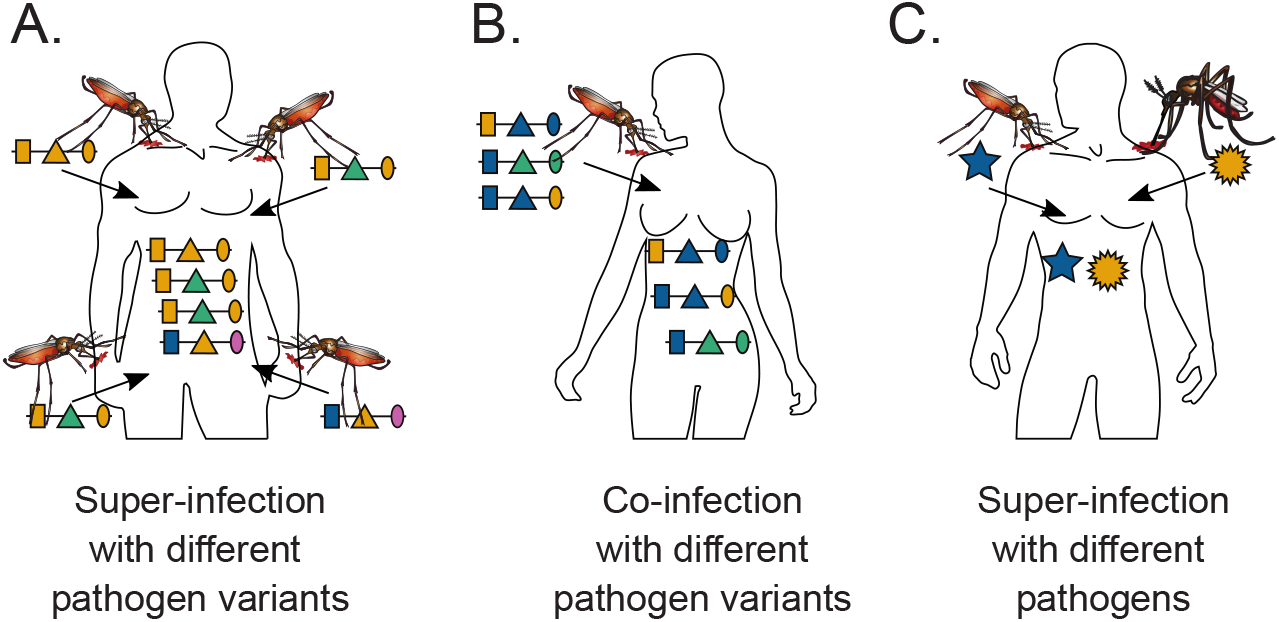
Super- and co-infections: Illustrated is the difference between super- and co-infections in the case of vector-borne diseases. **(A)** shows 4 super-infections (MOI = 4) with pathogenic variants, i.e., four independent infective events. At each infective event, one pathogenic variant is transmitted. Pathogenic variants are characterized genetically by their allelic expressions (colors) at three positions (shapes) in the genome, which is illustrated by the horizontal lines. Note that MOI = 4, although only three distinct haplotypes are transmitted because two vectors transmit the same pathogenic variant. **(B)** illustrates a co-infection with three pathogenic variants, i.e., a single infective event at which three pathogenic variants are transmitted. **(C)** illustrates a super-infection with two different pathogens, illustrated by different shapes, transmitted by different vector species.

‘In what follows, the terms “pathogens variants”, “lineages”, and “haplotypes” are used synonymously.

### Statistical model

#### Genetic architecture

Pathogen haplotypes are characterized by their allelic configuration at *L* marker loci. Haplotypes are denoted by vectors ***h*** = (*h*_1_, …, *h*_*L*_), where *h*_*k*_ represents one of the *n*_*k*_ possible alleles at locus *k*. This yields a total of 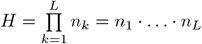 possible haplotypes. The set of all possible haplotypes is given by 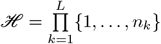 (see illustration Fig. 8A).

**Fig 8.**
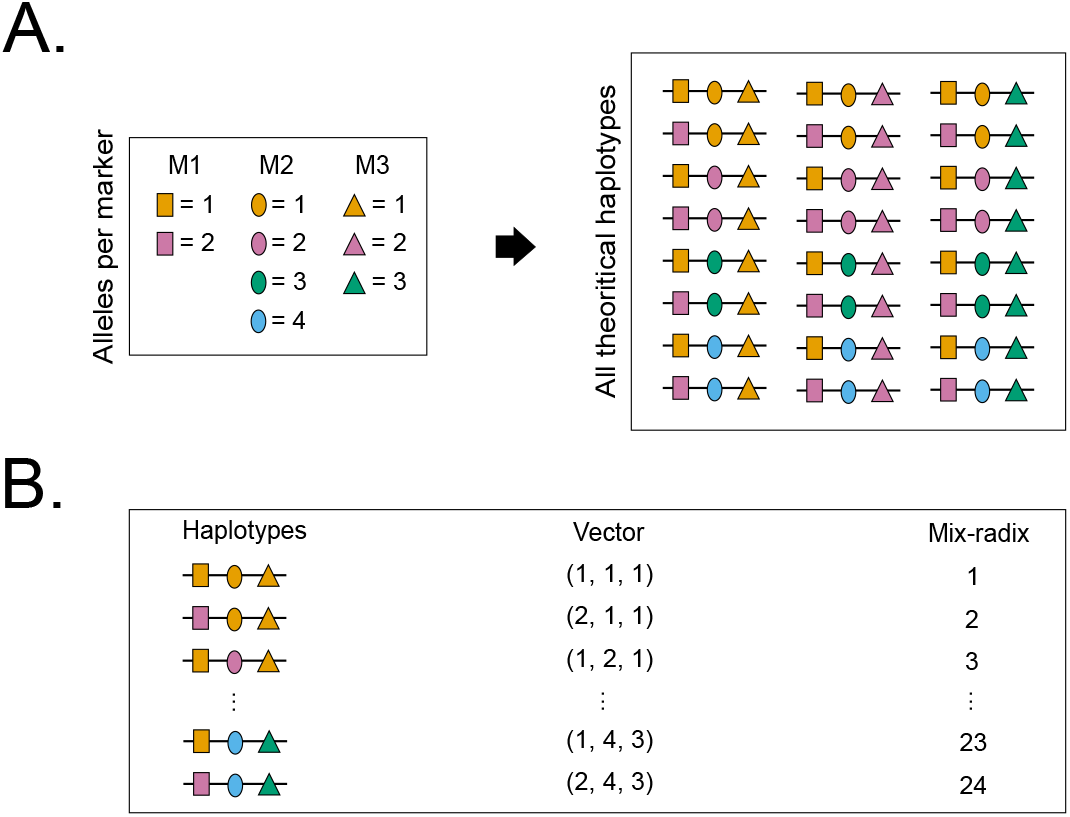
Genetic architecture and haplotype ranking: Panel **(A)** shows for a given genetic architecture (i.e., three loci with two, four, and three alleles at the first, second, and third locus, respectively), all the haplotypes that could be present in the pathogen population. Each shape represents a locus and each color is an allele at a given locus. All the possible haplotypes alongside their vector and mix-radix representations, i.e., ranking, are shown in panel **(B)**.

In the following, it will be convenient to label (rank) haplotypes by the numbers 1, …, *H*. The rank of hapotype ***h*** is denoted by *r*(***h***). A natural way of ranking is to assume that ***h*** = (*h*_1_, …, *h*_*L*_) is the mixed radix-representation [43] of the rank *r*(***h***) with base (*n*_1_, …, *n*_*L*_) (see Fig. 8B). More precisely,

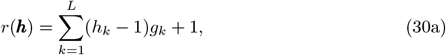

with

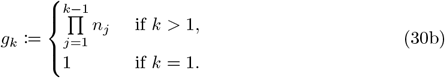

Ranking haplotypes has the advantage that each haplotype ***h*** can be uniquely identified with its rank *r*(***h***). While this is not necessary for the model derivation, it is the basis of an efficient model implementation.

The relative abundance of haplotype ***h*** in the pathogen’s population is denoted by *p*_***h***_, or *p*_*k*_ = *p*_***h***_ if ***h*** is the *k*-th haplotype, i.e., if *k* = *r*(***h***). Collectively, the vector of haplotype frequencies is denoted by

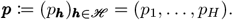

This notation implies that the set of haplotypes is regarded as ordered by the haplotype ranks.

Importantly, the number *H* of possible haplotypes increases “geometrically” with the number of loci *L* and alleles *n*_*k*_, *k* ∈ 1, …, *L*. However, only a subset of possible haplotypes will be realized in the pathogen population, i.e., *p*_***h***_ = 0 for some (or if *L* large most) haplotypes. It is assumed that the genetic architecture already subsumes all possible haplotypes so that mutational events do not need to be considered. In malaria, the pathogen population (or rather its reference point) is the population of sporozoites within the mosquitoes’ salivary glands [9, 10].

#### Distribution of MOI

MOI, which is sometimes also referred to as complexity of infection (COI) [4, 44, 45], is defined here as the number of independent infectious events during one disease episode assuming that exactly one pathogen variant (haplotype) is transmitted at each event (see Fig. 9). Notably, MOI does not refer to the number of distinct pathogen variants within an infection but to the number of infectious events, i.e., a host can be infected with the same pathogen variant multiple times (Fig. 9). Because it cannot be decided in practice how often a given pathogen variant infected a host, MOI is an unobservable quantity. Note that this definition of MOI coincides with the one from [8, 9] (cf. [2, 21] for more discussion).

**Fig 9.**
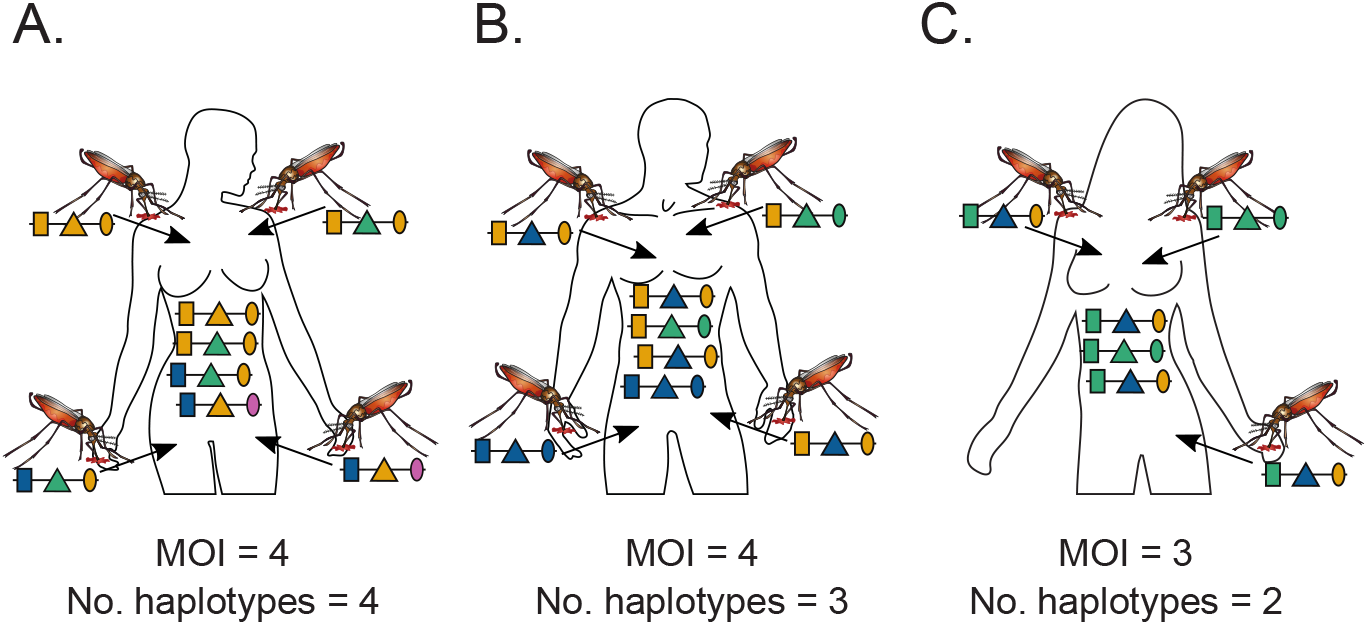
MOI and number of haplotypes: Illustrated are three hypothetical infections with pathogenic variants. Panel A shows four super-infections (cf. Fig. 7A, i.e., MOI = 4, with four different haplotypes. Panel B shows four super-infections (MOI = 4) with three different haplotypes. Panel C illustrates three super-infections (MOI = 3) with two different pathogenic variants.

Let the distribution of MOI on the population level be denoted by

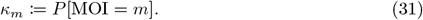

In the following only disease-positive individuals are considered, hence MOI≥ 1.

The intuitive assumption is that infective events are rare and independent. This implies that MOI follows a conditional Poisson distribution, i.e.,

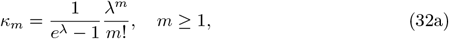

where *λ >* 0 is the parameter characterizing the distribution. Clearly, other distributions can be assumed. In malaria, for instance, exposure to mosquitoes might be heterogeneous across different groups in the population. Even if MOI is Poisson distributed in each group, in the total population MOI would then correspond to a mixture of Poisson distributions. In the limit case of infinitely many population strata, such that the Poisson parameter follows a beta distribution across the strata, MOI would follow a conditional negative binomial distribution. The negative binomial distribution at first might seem more appropriate than the conditional Poisson distribution, as it fits empirical observations of mosquito biting behavior better (cf. [46]). However, as noted in [19], maximum likelihood estimation of the negative binomial distribution is problematic (cf. also [47]) as the maximum-likelihood estimate does not exist if the empirical observations are not overdispersed. (Note also that a negative binomially distributed biting rate, does not necessarily imply that the number of infective events are negatively binomially distributed – in fact, these might be well approximated by the Poisson distribution). As an alternative, [19] explored a non-parametric MOI distribution (in the simple case of one marker locus, *L* = 1). This is the most flexible assumption. Their results suggest that even for rather overdispersed true MOI distributions, a statistical model based on the Poisson distribution performs similarly to their non-parametric alternative. As a consequence, in what follows, it is assumed that MOI follows a conditional Poisson distribution.

The mean of the conditional Poisson distribution is given by

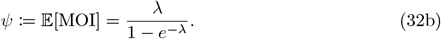

This parameter also characterizes the conditional Poisson distribution and is more intuitive than *λ*, as it is the average number of times infected individuals are super-infected.

#### Infections

Since a host can be super-infected multiple times with the same haplotype, an infection corresponds to the MOI vector ***m*** := (*m*_***h***_)_***h***∈*ℋ*_, where *m*_***h***_ is the number of times the host was infected with haplotype ***h***. Because only one haplotype is transmitted per infectious event, the MOI value of infection ***m*** is 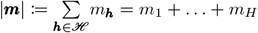.

Super-infections are assumed to be independent, i.e., given a host is infected with the pathogen, the host is not more or less likely to be infected again during the same disease episode. As a consequence, the probability of observing infection ***m***, given it has MOI |***m***| = *m*, follows a multinomial distribution with parameters *m* and ***p***, i.e.,***m***|*m*∼Multi(*m*,***p***). Hence,

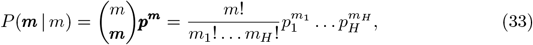

i.e., *m* haplotypes are drawn with replacement from the pathogen population according to their relative abundances ***p***.

#### Observations

As mentioned above, MOI is an unobservable quantity, because an infection ***m*** is itself unobservable, as it cannot be distinguished how often each haplotype was infecting. At most, the absence or presence of haplotypes can be observed, i.e., at most sign *m*_***h***_ is observable. However, in practice, pathogenic variants within an infection are determined by molecular assays. These typically generate unphased haplotype information, i.e., they generate consensus sequences that only distinguish the allelic variants at each marker locus, rather than determining the actual haplotypes present (see Fig. 10 for an illustration). In other words, at each marker locus, only the absence and presence of the alleles are observable.

The observable allelic information in a disease-positive sample is denoted by the vector ***x*** = (*x*_1_, …, *x*_*L*_), where *x*_*k*_ is the set of alleles detected at locus *k*, i.e.,

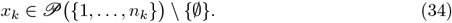

**Fig 10.**
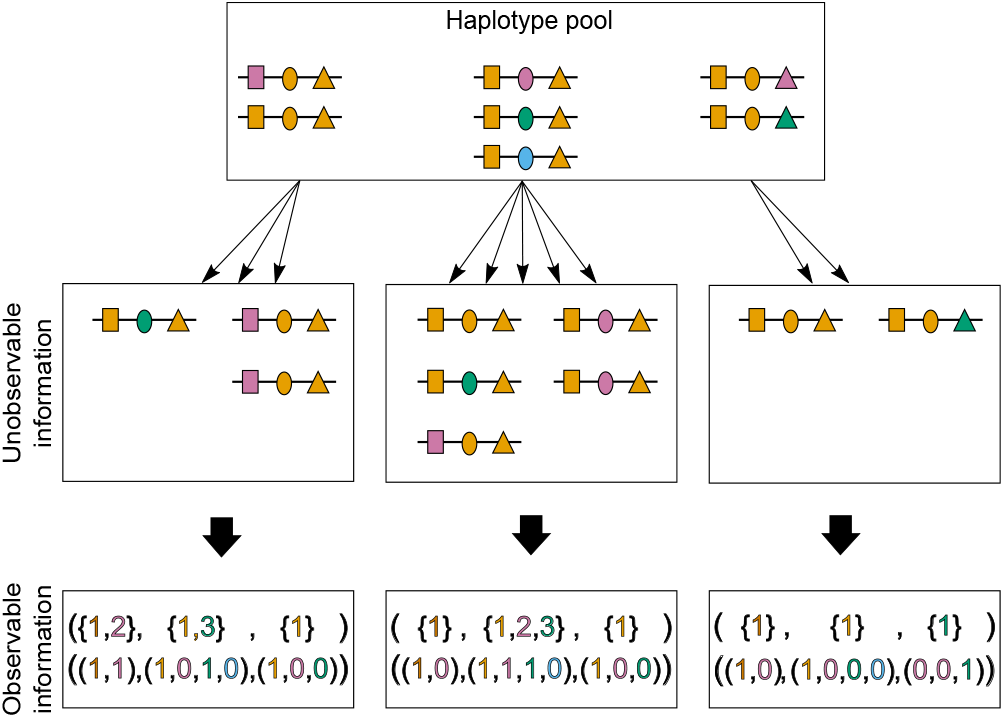
Haplotype information problem: Illustration shows three different infections from the same pathogen population. The first infection (middle left) describes a super-infection with two haplotypes, i.e., MOI = 2. The haplotype combination generates unphased haplotype information, i.e., ambiguous observation (bottom left). In this case, haplotype present in the observation cannot be reconstructed, as well as the corresponding MOI. The second infection (middle), illustrates a super-infection with four haplotypes, three of which are transmitted once while the fourth one is transmitted twice, i.e., MOI = 5. The resulting observation and MOI are ambiguous. The last infection (middle right) involves two haplotypes, i.e., MOI = 2, with resulting unphased haplotype information.

Here, *𝒫* denotes the power set. Hence, the components of ***x*** are sets (corresponding to 0–1 vectors of length *n*_*k*_; this correspondence is also illustrated in Fig. 1). Note, that *x*_*k*_ cannot be the empty set, is because we only consider disease-positive samples, and we assume the molecular assays to be free of errors (no missing or wrong information, cf. [22]). Therefore, a proper sample is one for which at least one allele is detected at each of the *L* loci. Consequently, the observational space is given by the set of all possible observations, i.e., as

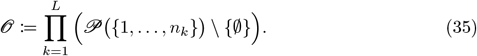

Note that, different infections ***m*** can yield the same observation ***x*** (see Fig. 11). Given MOI *m*, the set of all possible MOI vectors ***m*** vectors that lead to ***x*** is denoted by

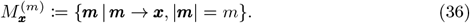

**Fig 11.**
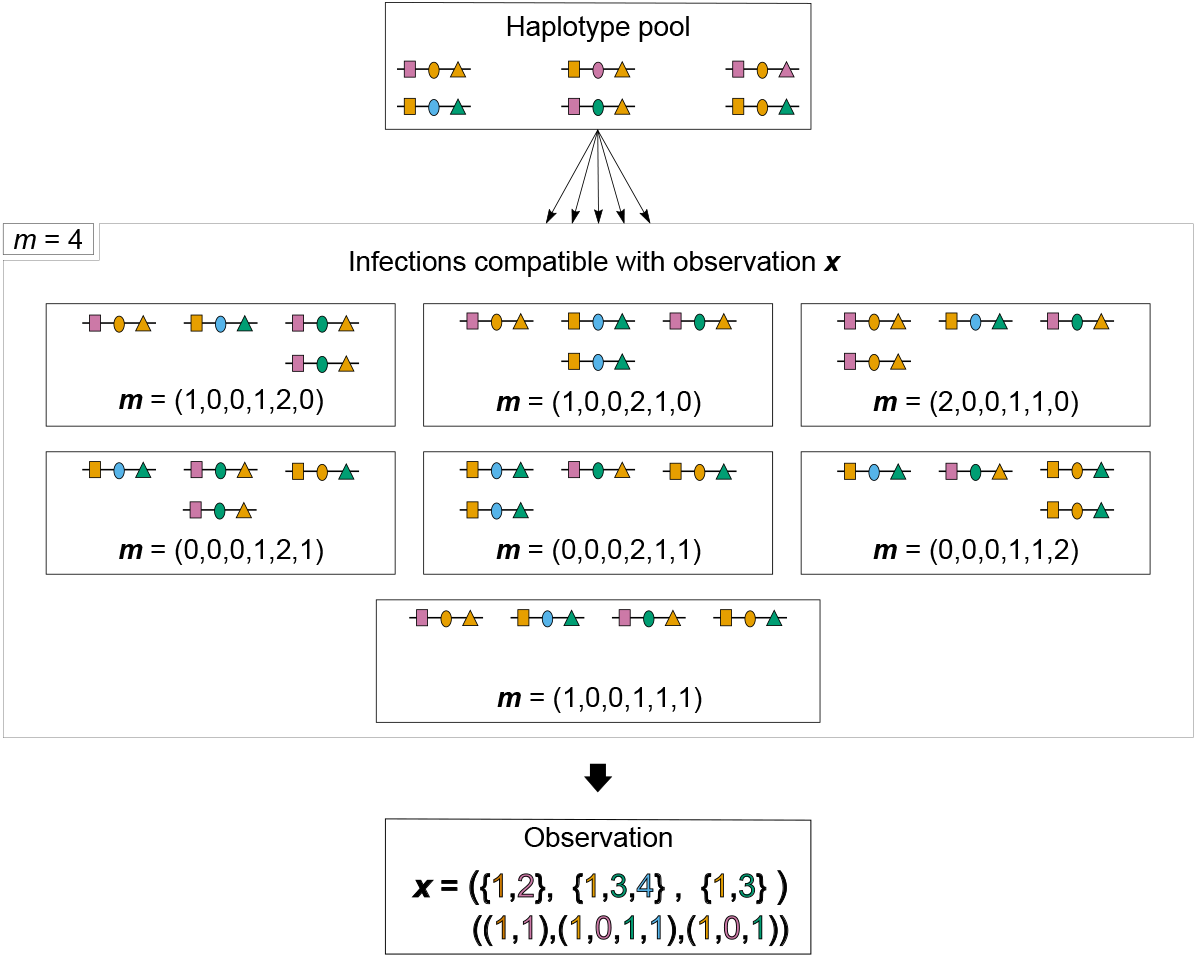
Compatibility between infections and observation: Illustration shows in the middle panel, for MOI = 4, all possible infections ***m*** = (*m*_1_, *m*_2_, *m*_3_, *m*_4_, *m*_5_, *m*_6_), i.e., |***m***| = *m* = 4 compatible with observation ***x*** (bottom panel). Haplotypes are sampled from a pool (top panel), in which they are ordered from one to six (so that the first and sixth haplotypes are at the top left and bottom right corner, respectively).

#### Distribution of observations

Two definitions are helpful to derive the probability distribution of the observations ***x*** ∈ *𝒪*. First, the set of all haplotypes which are compatible with observation ***x***, i.e.,

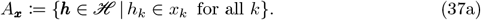

(When using the correspondence of *x*_*k*_ to a 0-1 vector, *h*_*k*_ ∈ *x*_*k*_ corresponds to *x*_*k,h*_*k* = 1, where *x*_*k,h*_*k* is the *h*_*k*_-th component of the 0-1 vector.) Hence, if ***h*** ∈ *A*_***x***_, haplotype ***h*** is potentially present in the infection underlying observation ***x***, while ***h∉*** *A*_***x***_ cannot be present. Second, the set of all sub-observations of ***x***, i.e.,

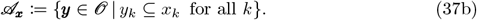

(When using the correspondence of *x*_*k*_ and *y*_*k*_ to a 0-1 vector, *y*_*k*_ ⊆ *x*_*k*_ corresponds to *x*_*k*_ ≤ *y*_*k*_, where the inequality is understood componentwise.) Hence, for a sub-observation ***y*** in *𝒜*_***x***_, any haplotype ***h*** that is compatible with observation ***y*** is also compatible with observation ***x***, i.e., if ***y*** ∈ *𝒜*_***x***_, then ***h*** ∈ *A*_***y***_ ⇒ ***h*** ∈ *A*_***x***_.

The probability of observation ***x*** given MOI= *m >* 0 is

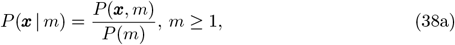

where *P* (*x, m*) is the probability of ***x*** descending from an infection with MOI= *m >* 0 and *P* (*m*) = *κ*_*m*_ is the probability of MOI= *m*. Recall that for ***m*** in 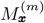, ***m*** yields ***x***. Hence, the law of total probability gives

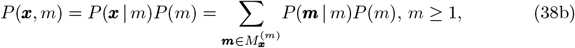

and

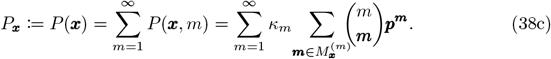

The probability mass function in (38c) can be further manipulated using the approach described in [12]. Namely, the partial order ⪯

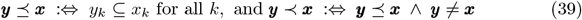

is defined on the set *𝒪*. (Since the components of an observation ***x*** correspond to 0-1 vectors *y*_*k*_ ⊆ *x*_*k*_ corresponds to *y*_*k*_ ≤ *x*_*k*_, where the inequality is understood componentwise.) Note, ***y*** ⪯ ***x*** is equivalent to ***y*** ∈ *𝒜*_***x***_. The set 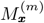 can be expressed as

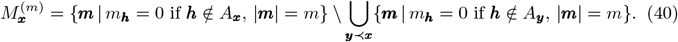

Here, infections ***m*** in the first set {***m***| *m*_***h***_ = 0 if ***h*** *∉ A*_***x***_, |***m***| = *m*} yield either observation ***x*** or a proper sub-observation ***y*** ≺ ***x***. The MOI vectors that correspond to such proper sub-observations ***y*** are removed by the second term.

Using the above, the inner-sum in (38c) can be rewritten using the inclusion-exclusion principle as

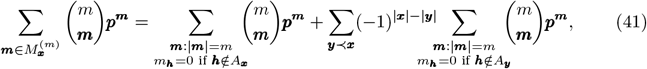

where 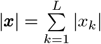 and 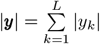 are respectively, the cardinals of ***x***, and ***y***. Note that 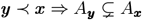 and |***x***| − |***y***| *>* 0. Hence,

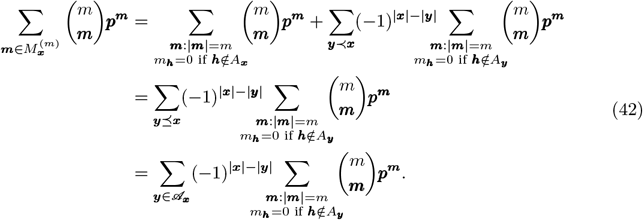

Therefore, the probability mass function in (38c) becomes

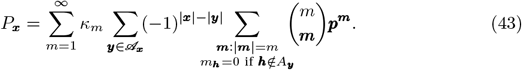

By the multinomial theorem, we have

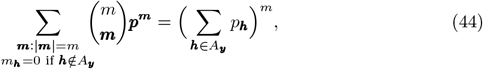

which can be replaced in (43) to give

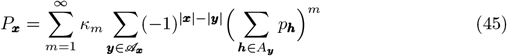

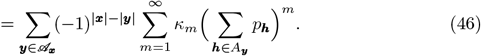

The inner-sum in (46) is the probability-generating function (PGF) of the MOI distribution, evaluated at 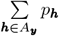. We denote the probability-generating function by *G* such that

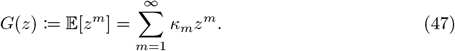

Therefore, the probability of observing ***x*** is given by

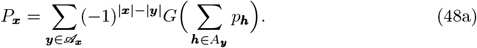

In the case that MOI follows a conditional Poisson distribution (as assumed here),

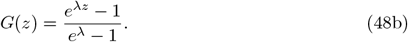

Under this assumption, the parameter space of the model is

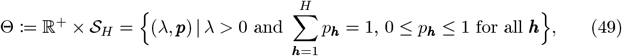

where 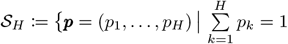 and 0 ≤ *p*_*k*_ ≤ 1, for all *k*} is an (*H* − 1)-dimensional simplex, and *λ* is a positive real number.

### Data sets and likelihood function

A typical molecular data set *𝒳* consists of *N* samples in the observational space *𝒪* (35). *I*.*e*., *N* set-valued vectors with *L* components corresponding to alleles detected at each of the *L* markers. The *j*-th sample is denoted by using a superscript, i.e., 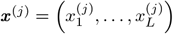, where 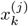 is the allelic information at locus *k*.

Let *n*_***x***_ be the number of times each observation ***x*** is made in the data set. Hence,

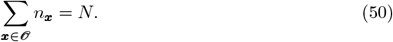

Consequently, the data set *𝒳* can be represented as the integer-valued vector

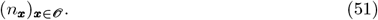

This representation will be repeatedly used in the following derivations. Note that, for increased number of loci *L* and number of alleles *n*_*k*_, not all possible observations are detected, i.e., 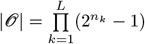 exceeds the sample size *N*. Therefore, *n*_***x***_ = 0 for most observations ***x*** ∈ *𝒪*.

Given *𝒳*, the model parameters ***θ*** := (*λ*, ***p***) Θ can be collectively estimated by maximum likelihood. Because the *N* samples are assumed to be independent and identically distributed, the likelihood function ℒ_*𝒳*_ (***θ***) is

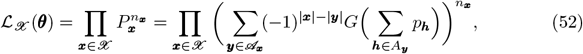

and the log-likelihood function becomes

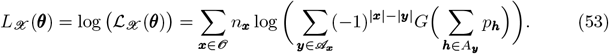

The maximum likelihood estimate (MLE) 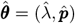 of the model parameters is obtained by maximizing the log-likelihood function. A closed solution for the MLE is typically not possible, particularly for this complex likelihood function. In fact, a closed solution is not even possible in the simplest case of a single molecular marker (cf. [9, 22]).

Here, the expectation-maximization (EM)-algorithm is used to derive the MLE. It is an efficient, stable, and fast-converging algorithm to maximize the likelihood function [24]. The algorithm is derived in section Maximum likelihood estimate.

Because the MLE needs to be derived numerically, it is impossible to obtain analytic results about the quality of the estimator. Particularly, bias and variance cannot be determined explicitly. However, these can be assessed numerically as it was done for the special cases of (i) a single molecular marker (*L* = 1) [25], (ii) *L* biallelic markers [12], and (iii) two multiallelic markers (*L* = 2) [10]. The results suggested that the estimator is asymptotically unbiased, and its asymptotic variance is given by the Cramér-Rao lower bound (CRLB). This is confirmed by asymptotic results in the special case of a single marker (*L* = 1), in which case, the model falls into the exponential-family class. Given these facts, the present estimator can be considered asymptotically unbiased and efficient. However, for the asymptotic variance, the CRLB has to be derived for the present general case.

### Numerical investigation of finite sample properties

Finite sample properties of the estimator are investigated by numerical simulations.

#### data set generation

Data sets are created following Fig. 10. Specifically, for a choice of the parameters *λ* and ***p***, a data set *𝒳* of size *N* is created by first sampling *N* MOI values (*m*^(*i*)^, *i* = 1, …, *N*) according to a conditional Poisson distribution. Next, *N* MOI vectors ***m***^(*i*)^ (*i* = 1, …, *N*) are drawn from multinomial distributions with parameters *m*^(*i*)^ and ***p***. From each MOI vector ***m***^(*i*)^, the observation ***x***^(*i*)^ is created, by retaining only the information concerning the absence and presence of alleles at each marker.

For each set of parameters (*λ, p, N*), this procedure is repeated *K* = 100 000 times to generate *K* data sets *𝒳* _1_, …, *𝒳* _*K*_ of sample size *N*.

#### Simulated variance

Given a set of parameters *λ*, ***p***, and *N*, for each of the *K* data sets *𝒳* _*k*_, the MLEs 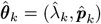 are calculated and the estimate of the mean MOI 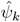 is derived. Next, the variance of the estimator for a parameter of interest, *θ* = *ψ, p*_1_, …, *p*_*H*_, is calculated as the empirical variance, i.e.,

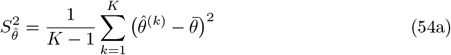

where

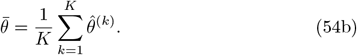

To allow comparison between different true values of the MOI parameter *λ*, the variation of the mean MOI parameter is reported as the coefficient of variation (CV), i.e.,

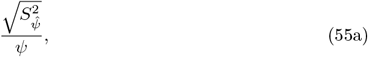

whereas the variation of the haplotype frequencies is reported in terms of the empirical standard deviation 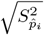. These quantities are compared to the appropriate transforms of the CRLB, i.e., to 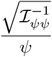 and 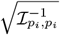.

#### Parameter choices for numerical investigations

The following parameters are chosen for numerical investigations.

##### Genetic architecture

Two markers (*n* = 2) are assumed. Moreover, for the simulations, *n*_1_ = *n*_2_ = 2 and *n*_1_ = 4, *n*_2_ = 7 loci are chosen, leading respectively to *H* = 4 and *H* = 28 possible haplotypes.

##### MOI parameter

The choice of MOI parameters is made to represent different levels of transmission intensities. With malaria in mind, MOI parameters are chosen as *λ* = 0.1, 0.25, 0.5, 1, 1.5, 2, 2.5, corresponding to mean MOI *ψ* = 1.05, 1.13, 1.27, 1.58, 1.93, 2.31, 2.72. Note that *ψ <* 1.27 corresponds to low transmission, 1.27 ≤ *ψ <* 1.93 to intermediate transmission and *ψ* ≥ 1.93 to high transmission [21].

##### Haplotype frequency distribution

The distribution of haplotype frequency ***p*** is chosen to mimic two extreme cases:

(i) a uniform (balanced) distribution such that

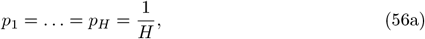

and (ii) an unbalanced distribution for which one haplotype, e.g., *p*_1_ is predominant with a frequency of 70%, while the remaining haplotypes have the same frequency, i.e.,

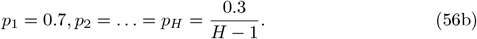

In particular, for the chosen genetic architectures, this gives *p*_1_ = 0.7, *p*_2_ = *p*_3_ = *p*_4_ = 0.1 in the case *n*_1_ = *n*_2_ = 2 and 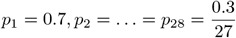 in the case *n*_1_ = 4 and *n*_2_ = 7.

##### Sample size

Sample sizes *N* = 50, 100, 150, 200, 500 are chosen for the simulations. Note, in practice, sample size correlates with transmission intensity. In areas of low disease transmission, it is more challenging to achieve a large sample size, whereas this is much easier in areas of high transmission.

### Population genetic measures

In a population of size *N* the expected heterozygosity 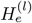 at locus *l* with *n*_*l*_ segregating alleles 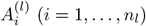, each with frequency 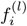, is given by

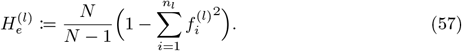

Note that 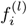 is obtained by marginalization of the frequencies of haplotypes containing allele 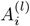 at locus *l*, i.e.,

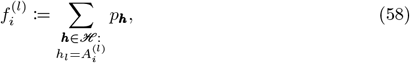

the sum of the frequencies of all haplotypes, which carry allele 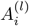 at locus *l*. Consider a subset of loci *S* ⊆ {1, …, *L*} and a specific allelic configuration 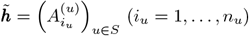, i.e., a “sub-haplotype” determined by its allelic configuration at the markers in set *S*. One can calculate the heterozygosity at locus *l*, conditioned on the background of the sub-haplotype 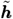. The marginal frequency of allele 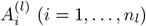 at locus *l* is given by

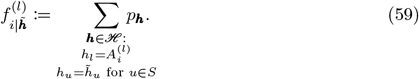

The heterozygosity conditioned on 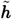 is then given by

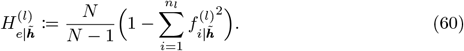

Pairwise LD can be obtained from several metrics (cf. [48, 49]). Here, with no particular preference, two of these commonly used metrics, i.e., *r*^2^ (also known as *D*^*^) and *D*^′^ are reported. The LD metric 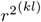 between two loci *k* and *l* is obtained based on the Hardy-Weinberg heterozygosities 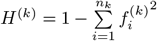 and 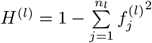 at loci *k* and *l*, respectively, as

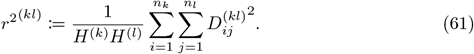

where 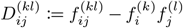 measures the discrepancy between the observed frequency 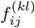 of sub-haplotype 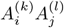 and its expected frequency 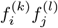 under random association.

The measure *D*^′(*kl*)^ is defined in [48, 49] as

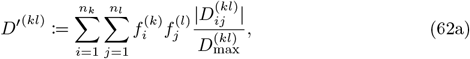

where

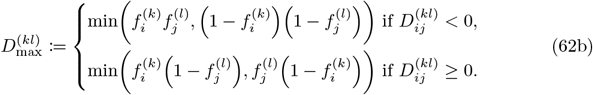

Restricted to the background of a given sub-haplotype 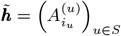, the LD measures are based on conditional allele frequencies defined as in (59). The conditional LD measures 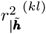 and 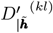 are defined by

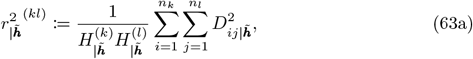

with

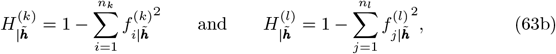

and

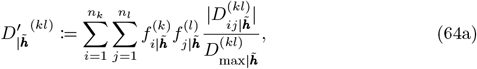

with

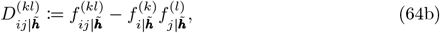

and

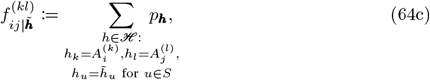

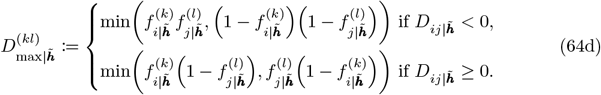

## Acknowledgments

The authors gratefully acknowledge the African Institute for Mathematical Sciences (AIMS) Cameroon to have favored the meeting between the authors and supporting this research. The many fruitful discussion with friends and colleagues that helped to improve the manuscript were highly appreciated. This research is supported by a grant from the German Academic Exchange (“Mathematics against malaria within the AIMS network”, project-ID 57417782), DFG (“Ökologisch nachhaltige Wertschöpfungsketten in der Landwirtschaft durch Optimierung des Insektizid-Gebrauchs aufgrund von automatisiertem Schädlings-Monitoring”) and the SMWK (“Vorlaufforschung Technologieentwicklung 4.0”). The content is solely the responsibility of the authors and does not represent the official views of anybody.

## Supporting information

**S1 Appendix. Mathematical Appendix**

**S2 Appendix. R-script manual**

**S1 R Scripts and example data sets. Zip file containing an implementation of the model as an R script, a template R script with commands to use for data analysis using the method, and an example molecular dataset**.

